# Mechanisms determining *Schistosoma mansoni* CRAC channel activation

**DOI:** 10.64898/2026.08.03.742424

**Authors:** Ana Eliza Zeraik, Olivier Romito, Aparna Gudlur, Kenneth A. Stauderman, Gönül Veliçelebi, Ana Paula Ulian Araujo, Mohamed Trebak, Patrick G Hogan

**Affiliations:** Division of Signalling and Gene Expression, La Jolla Institute for Immunology, La Jolla, CA 92037, USA; Instituto de Física de São Carlos, Universidade de São Paulo, São Carlos 13563-120 SP, Brazil; Laboratório de Química e Função de Proteínas e Peptídeos, Centro de Biociências e Biotecnologia, Universidade Estadual do Norte Fluminense Darcy Ribeiro, Campos dos Goytacazes, Rio de Janeiro 28013-602, Brazil; Department of Pharmacology and Chemical Biology, University of Pittsburgh School of Medicine, Pittsburgh, PA 15213, USA; CalciMedica, Inc., 505 Coast Blvd. South, Suite 306A, La Jolla, CA 92037, USA; Camino Pharma, 9920 Pacific Heights Blvd, Suite 150 San Diego, CA 92121, USA; Vascular Medicine Institute, University of Pittsburgh School of Medicine, Pittsburgh, PA 15213, USA; UPMC Hillman Cancer Center, University of Pittsburgh School of Medicine, Pittsburgh, PA 15213, USA; Program in Immunology, University of California–San Diego, La Jolla, CA 92093, USA; Moores Cancer Center, University of California–San Diego, La Jolla, CA 92037, USA

**Keywords:** Calcium signaling, schistosome, schistosomiasis, STIM, ORAI

## Abstract

*Schistosoma mansoni* and its schistosome relatives are parasitic worms that impose a substantial disease burden on human populations and livestock. On the rationale that calcium signalling is a critical process in multicellular organisms, we have examined wildtype and engineered *S. mansoni* STIM and ORAI— orthologues of STIM and ORAI known in mammals and other species for their central role in cellular calcium signalling— by imaging their localization, interactions, and contribution to ion currents and calcium influx in living cells. The ER membrane protein *S. mansoni* STIM recapitulates the essential functions of mammalian STIM1, namely, calcium-sensing by its ER-luminal domain, targeting to ER-plasma membrane junctions through interactions with the plasma membrane and with plasma membrane *S. mansoni* ORAI channels, and an ability to gate the *S. mansoni* ORAI channel. *S. mansoni* ORAI is a plasma membrane calcium channel that exhibits striking parallels with mammalian ORAI1 in its pore architecture and gating mechanism. The schistosome and human proteins are not completely interchangeable, however, and schistosome-human ORAI chimeras point to a special role of the ORAI N terminus in channel gating. Importantly, we demonstrate pharmacological differences between the schistosome and human channels that may offer an opportunity for selective therapeutic targeting of schistosome STIM-ORAI-dependent calcium entry.

**Author Summary:** Calcium channels represent potential targets to parasitic helminths. We investigated *Schistosoma mansoni* CRAC channel activation through the expression of its proteins. We have established that the fundamental protein conformational changes and protein-protein interactions underlying STIM-ORAI signaling are shared between humans and schistosome proteins. Importantly, a key finding is that evolutionary divergence in residues that are not implicated in the basic mechanisms of STIM-ORAI activation appears to offer a window for pharmacological inhibitors that would be selective for the schistosome ORAI channel. We identified pharmacological differences for two compounds tested. These differences open avenues for the development of selective drugs that can target the *S. mansoni* CRAC channel without affecting human physiology, thus offering the prospect of new treatments for schistosomiasis.

## Introduction

Schistosomiasis— caused by *Schistosoma mansoni*, *S. haematobium*, *S. japonicum*, and less prevalent species— is endemic in sub-Saharan Africa, the Middle East, Caribbean islands, parts of South America, and locales in Southeast Asia, and imposes a substantial health burden on rural and economically disadvantaged human populations (1–3). Other schistosome species that infect cattle, sheep, and goats can have severe economic and health impacts on some of these same communities (4–6). Current therapy directed at schistosomiasis remains dependent on a single medication, praziquantel, both for preventive programs of mass drug administration and for treatment (7, 8). Due to praziquantel side effects (9), lower efficacy with repeated treatment (10), inefficiency against early stages of *Schistosoma* species (11, 12) and risk of potential resistance (13, 14), an expanded pharmacological arsenal is desirable, especially if integrated ‘One Health’ programs aimed at controlling disease in both human and livestock populations are implemented.

The search for novel effective treatments for schistosomiasis is ongoing (15). In parallel, the *S. mansoni*, *S. japonicum*, and *S. haematobium* genomic sequences have been completed (16–21). Functional genomics in schistosomes is still challenging (22) but progress has been made, including genome editing using CRISPR-Cas9 (23–25) and single cell RNA-seq (26). A newly developed methodology for RNAi screening in schistosomes might identify new therapeutic targets (27). These approaches can be complemented by targeted analysis of key cellular signalling proteins.

Calcium signalling has critical roles in the development and physiology of eukaryotes, contributing both to broadly shared processes and to specialized roles shaped during evolution (28, 29). One key mechanism of cellular calcium signalling is store-dependent calcium entry (30–33). Its dependence on STIM proteins that serve as ER calcium sensors and ORAI proteins that form plasma-membrane calcium channels was initially established in studies of human, mouse, and *Drosophila* cells (34–38). The single STIM homolog and the single ORAI homolog in *S. mansoni*— here denoted *Sm*STIM and *Sm*ORAI— exhibit ample sequence conservation relative to the corresponding human proteins (Fig. S1), indicating that a close comparison of calcium signalling by the schistosome and human proteins might define the functional similarities and functional differences.

Transcripts of *Sm*ORAI are enriched in gonads of adult worms, pointing to a role in reproduction (39), but the mechanism and functional relevance of CRAC channel gating remains largely unknown in these parasitic worms.

The objectives of this work were to determine whether the basic mechanisms of STIM-ORAI signalling that have been defined for the human proteins are conserved in *S. mansoni*, and to explore whether there are sufficient differences in detail that selective pharmacological targeting is likely to succeed. Since schistosome primary cells and cell lines are not available (40), the initial experiments have been conducted by expressing *Sm*STIM and *Sm*ORAI in human cells. We have focused on essential conserved conformational changes and interactions, STIM-ORAI pharmacology, and observed features that distinguish the schistosome and human proteins.

## Results

### *Sm*ORAI is a STIM-gated calcium channel

mCherry-*Sm*ORAI was localized to the plasma membrane when expressed in HEK293 cells, was more or less uniformly distributed in the cell footprint, and was quiescent as a channel, as documented below. Co-expression of EGFP-*Sm*STIM with mCherry-*Sm*ORAI resulted in constitutive partial recruitment of the channels to presumed junctional sites marked by *Sm*STIM, although a substantial amount of *Sm*STIM and *Sm*ORAI remained outside junctions (Fig. 1A). Co-expression of *Sm*STIM also resulted in *Sm*ORAI channel activation measured as calcium influx (Fig. 1B). Thus, the SmSTIM–SmORAI combination in HEK293 cells generated a channel that was constitutively active, as shown by enhanced calcium entry upon restoration of extracellular calcium (Fig. 1B). Subsequent depletion of ER calcium stores with thapsigargin did not cause any further increase in calcium influx, indicating that the channel operates independently of store depletion. This is further supported by the minimal change or lack of a detectable change in *Sm*ORAI junctional localization. It may be that store-independent signalling reflects a missing restraint on *Sm*STIM-*Sm*ORAI activation in human cells, compared to the native environment in schistosome cells, but this explanation cannot be verified in the absence of a suitable schistosome cell preparation.

**Fig. 1.**
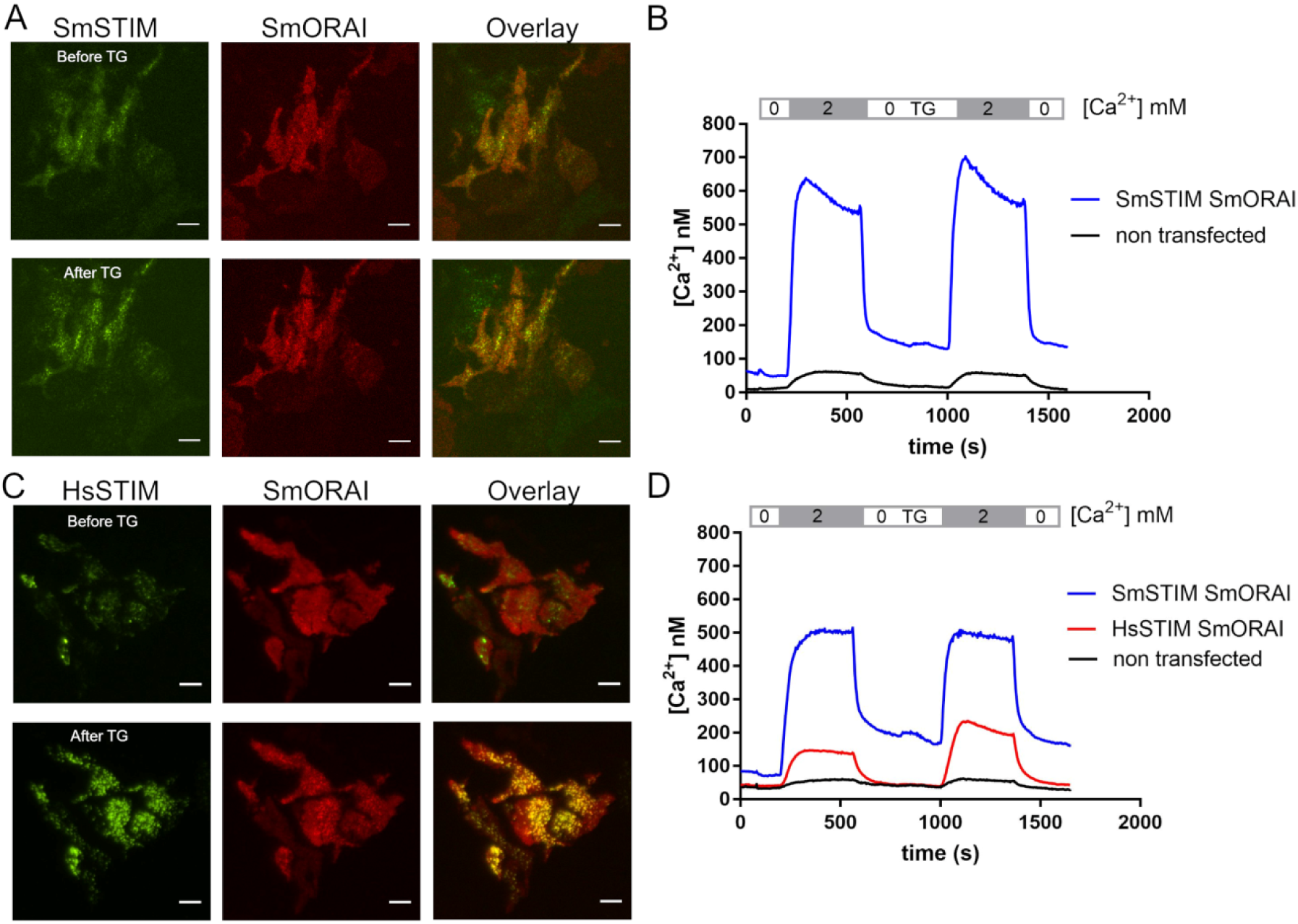
*Sm*Orai is a STIM-gated calcium channel. (A) TIRF images of HEK293 cells coexpressing EGFP-*Sm*STIM and mCherry-*Sm*Orai before (*upper panels*) and after (*lower panels*) store depletion with 1µM thapsigargin (TG). *Sm*STIM puncta are visible in the upper panels, before stimulation of the cells. Scale bar, 10 µM. (B) Cytoplasmic Ca^2+^ concentration monitored by Fura-2 fluorescence ratio in HEK293 cells coexpressing EGFP-*Sm*STIM and mCherry-*Sm*Orai (*blue*) and in non-transfected cells (*black*). (C) TIRF microscopy of HEK293 cells coexpressing EGFP-*Hs*STIM and mCherry-*Sm*Orai before (*upper panels*) and after (*lower panels*) stimulation with TG. Scale bar, 10 µM. (D) Cytoplasmic Ca^2+^ concentration in HEK293 cells coexpressing EGFP-*Hs*STIM and mCherry-*Sm*Orai (*red*) in comparison to cells coexpressing EGFP-*Sm*STIM and mCherry-*Sm*Orai (*blue*) and non-transfected cells (*black*). The basal activity of the channel in the red trace, upon switching from a nominally Ca^2+^-free conditions to 2 mM Ca^2+^, is probably due to modest overexpression of *Hs*STIM1. TIRF and Ca²⁺ imaging experiments were performed at least three independent times. The images shown in Fig. 1A and C are representative. The graphs in Fig. 1B and D show the mean of at least 20 cells from one representative experiment.

In contrast, co-expression of *Sm*ORAI with human STIM1 (*Hs*STIM1) did result in store-dependent recruitment and activation of the *Sm*ORAI channels. There was little *Hs*STIM1 at junctions in resting cells, and little colocalization of *Sm*ORAI with *Hs*STIM1 (Fig. 1C). Depleting ER calcium stores with thapsigargin elicited relocalization of *Hs*STIM1 to ER-PM junctions, as expected, together with an incomplete recruitment of *Sm*ORAI to junctions (Fig. 1C), and store-dependent calcium influx (Fig. 1D). The store-dependent calcium signal elicited by *Hs*STIM1 coexpressed with *Sm*ORAI was consistently less than the constitutive calcium signal observed with *Sm*STIM and *Sm*ORAI. This observation is in line with ORAI chimera experiments described below, which indicate that *Hs*STIM1 may be less effective than *Sm*STIM in activating the schistosome channels.

Further direct evidence that *Sm*ORAI is a STIM-gated channel came from its constitutive, store-independent activation by coexpressed *Sm*STIM and *Hs*STIM1 cytoplasmic fragments, *Sm*STIM(212-489) and *Hs*STIM1(233-473) respectively (Fig. S2). Constitutive channel activation by STIM1 cytoplasmic fragments was part of the early evidence that human ORAI1 is gated by STIM1 (41).

Consistent with the calcium influx data, an inwardly rectifying current developed over time in whole-cell patch-clamp recordings from cells coexpressing either *Sm*Orai and *Hs*STIM1 (Fig. S3 A and B) or *Hs*Orai1 and *Hs*STIM1 (Fig. S3 C and D). The current promoted by *Sm*Orai and *Sm*STIM is characterized below (Figs 2-5).

**Fig. 2.**
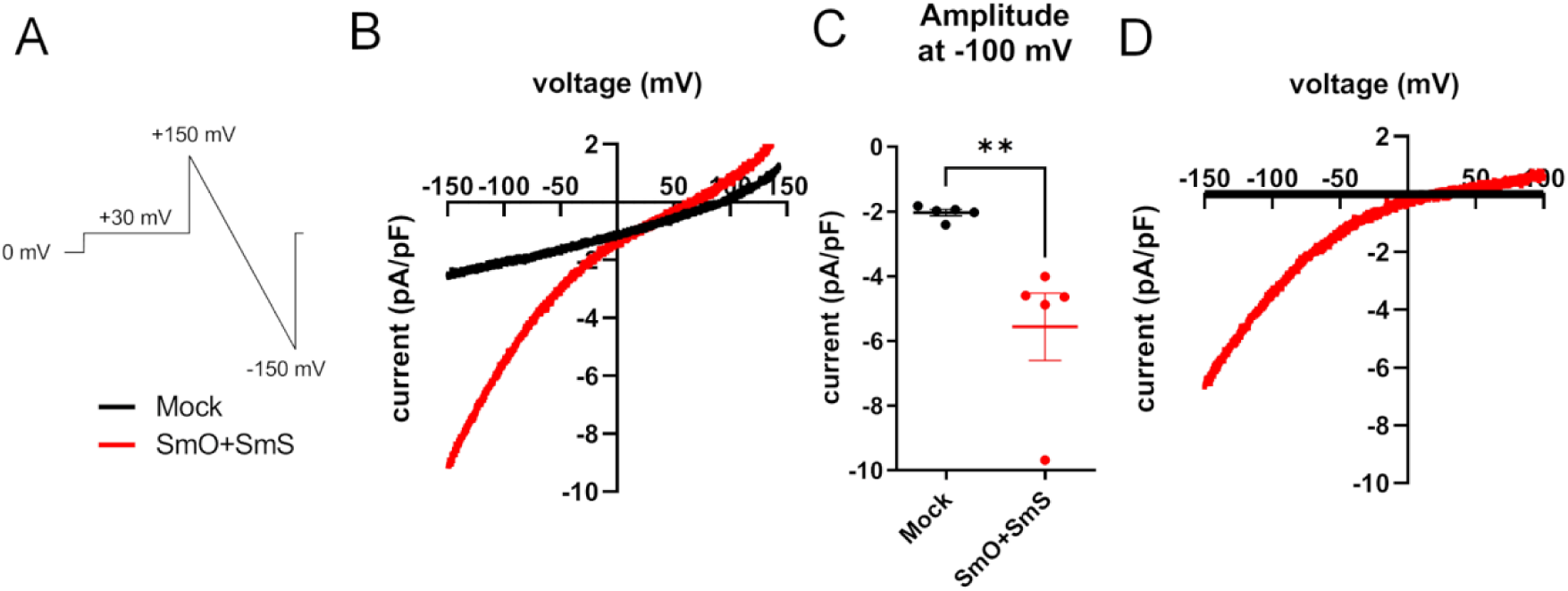
SmOrai/SmSTIM expression lead to a constitutively active current with inward rectification. (A) Protocol used for current recordings entails reverse voltage ramps from +150 mV to −150 mV, lasting for 250 ms, applied every 3s. (B) Average of I/V relationships from mock-transfected HEK-OraiTKO cells (N=5) and SmOrai/SmSTIM-expressing HEK-OraiTKO cells (N=5). (C) Current density at −100 mV from mock-transfected HEK-OraiTKO cells and SmOrai/SmSTIM-expressing HEK-OraiTKO cells. (D) Average I/V relationships from several recordings that are leak-subtracted from B.

### Electrophysiological characterization of the *Sm*ORAI channel

We coexpressed *Sm*STIM and *Sm*ORAI in human embryonic kidney 293 (HEK293) cells in which expression of all three endogenous human ORAI proteins had been eliminated by CRISPR/Cas9 targeting (ORAI-TKO cells) (42), and employed whole-cell patch clamp recording to determine the biophysical properties of *Sm*ORAI channel currents. We delivered a 250-ms reverse voltage ramp from +150 mV to −150 mV to minimize potential contamination with voltage-gated currents (Fig. 2A). In addition to the small leak current recorded in mock-transfected cells, cells coexpressing *Sm*STIM and *Sm*ORAI had a constitutive *Sm*ORAI current (Fig. 2B-C). After subtracting the leak current measured in mock-transfected cells, the *Sm*ORAI current exhibited strong inward rectification and an estimated reversal potential ranging from +6.0 mV to +86.9mV, average +49 ± 6 mV (*n*=13) (Fig. 2D).

Gd^3+^ at a concentration of 0.1-3 µM is sufficient to block native store-operated Ca^2+^ entry in wildtype DT40 B cells or wildtype HEK293 cells, by 90% or more, when Ca^2+^ entry is measured by Fura2 fluorescence in extracellular solutions containing 1.5-1.8 mM Ca^2+^ (^43^)^;^ and Gd^3+^ 5 µM is sufficient to block CRAC currents carried by human ORAI1, ORAI2, or ORAI3 coexpressed with human STIM1 in ORAI-TKO cells, when assayed in 20 mM external Ca^2+^ solutions (44). Surprisingly, 5 µM Gd^3+^ had no significant effect on *Sm*ORAI currents, and only 50 µM Gd^3+^ led to nearly complete inhibition of the currents (Fig. 3A-D). The estimated IC₅₀ for *Sm*ORAI channel inhibition by Gd^3+^ was 13 µM (Fig. 3D). Since lanthanides bind at the selectivity filter or the adjacent TM1-TM2 loops of the human and *Drosophila* CRAC channels (45–47), the comparatively lower efficacy of Gd^3+^ in this case implies a structural difference between the schistosome and human channels in the vicinity of the E50 selectivity filter.

**Fig. 3.**
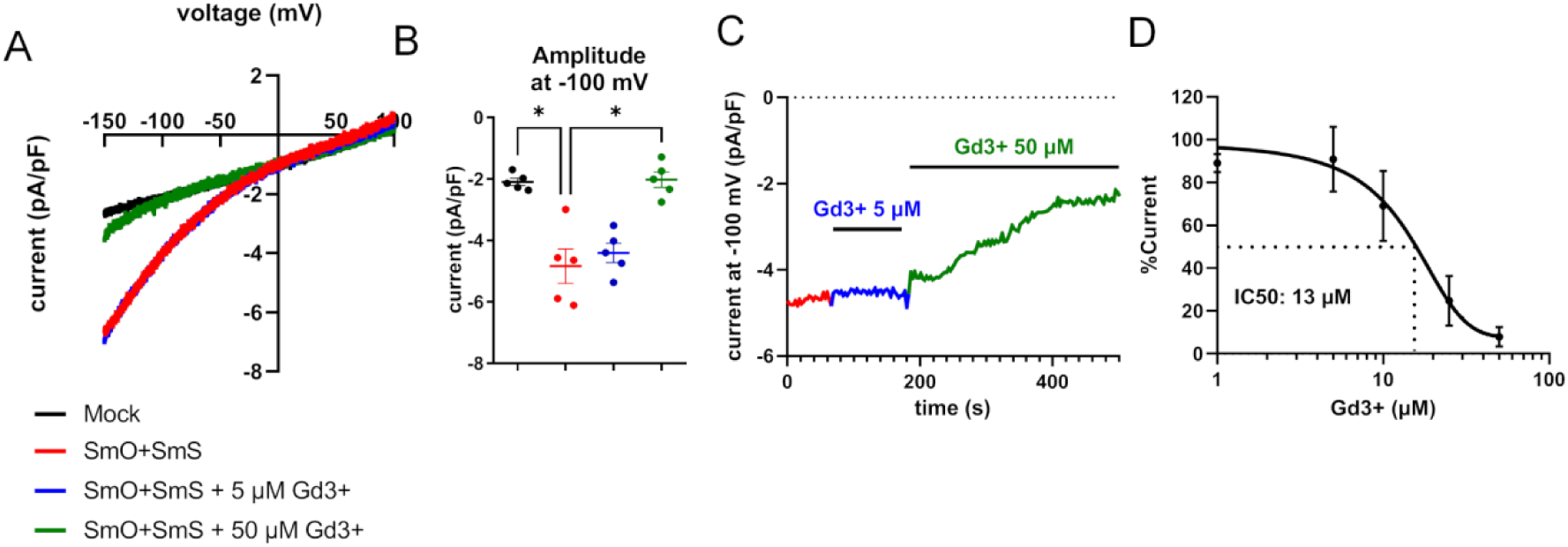
SmOrai/SmSTIM-mediated currents have lower sensitivity to inhibition by Gd^3+^. (A) Average of I/V relationships from mock-transfected HEK-OraiTKO cells and SmOrai/SmSTIM-expressing HEK-OraiTKO cells. SmOrai/SmSTIM-expressing HEK-OraiTKO cells were exposed to acute administration of either 5 μM or 50 μM (N=3). (B) Current density at −100 mV from mock-transfected HEK-OraiTKO cells and SmOrai/SmSTIM-expressing HEK-OraiTKO cells either untreated or treated with 5 μM or 50 μM Gd^3+^ (N=5). (C) Representative time course of whole cell current, measured at −100 mV in SmOrai/SmSTIM-expressing HEK-OraiTKO cells exposed to either 5 μM or 50 μM Gd^3+^.(D) Dose-response curve showing IC50 for Gd^3+^ on currents recorded from SmOrai/SmSTIM-expressing HEK-OraiTKO cells (Gd^3+^ 0 µM: N=9; 1 µM: N=4; 5 µM: N=5; 10 µM: N=4; 25 µM: N=3; 50 µM: N=5).

Next, we recorded *Sm*ORAI-mediated currents in a divalent-free (DVF) external solution. Mock-transfected cells showed only a small increase of the leak current at −100 mV in DVF solution (Fig. S4). In contrast, cells coexpressing *Sm*STIM and *Sm*ORAI exhibited a large inward Na^+^ current at −100 mV that depotentiated with time (Fig. S4 B), similarly to the depotentiation of human CRAC currents in DVF external solutions (48). Leak subtraction revealed an I-V curve dominated by the inward presumed Na^+^ current at negative potentials (Fig. S4 C,D).

Human CRAC currents undergo rapid calcium-dependent inactivation (CDI) due to local Ca^2+^ feedback when 10 mM of the slow Ca^2+^ chelator EGTA is included in the pipette solution, but not when the fast chelator BAPTA is used (49). To test for CDI in cells coexpressing *Sm*STIM and *Sm*ORAI, we applied 250-ms voltage steps over a range from −140 mV to +140 mV (Fig. S5), and compared the evoked currents when 10 mM EGTA was included in the patch pipette to currents recorded with 20 mM BAPTA in the pipette solution. CDI was not evident in either condition, indicating that, unlike human CRAC currents, schistosome CRAC currents do not undergo fast CDI (Fig. S5 B-K). The lack of fast CDI is a feature that schistosome CRAC currents share with *Drosophila* CRAC currents (50).

### *Sm*ORAI conserves channel pore architecture and gating mechanism

We next examined whether the *Sm*ORAI channel pore and the gating mechanism are similar to those of mammalian ORAI channels. Guided by the extensive mapping of mutations that alter the function of the human ORAI1 channel (51), we mutated critical residues in *Sm*ORAI transmembrane segment 1 (TM1) lining the channel pore, in TM2, in TM3, and in the cytoplasmic C-terminal region. The locations of mutations studied and the corresponding *Sm*ORAI and *Hs*ORAI1 residue numbering are indicated in Fig. 4 A; (see also the *Sm*ORAI-*Hs*ORAI1 sequence alignment in Fig. S1).

**Fig. 4.**
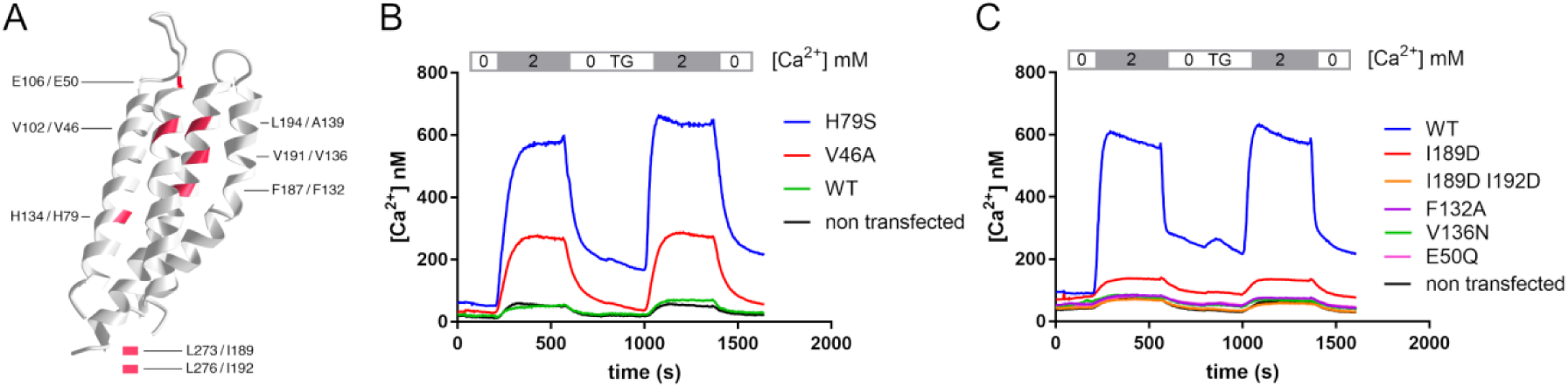
*Sm*Orai mutations. (A) Richardson diagram of a *Drosophila* Orai monomer (PDB ID 7KR5, chain E) (80) with pore-lining transmembrane helix 1 in the foreground, center. Sites of mutations are demarcated by red shading and by labels identifying the corresponding human / schistosome residues. Sites of mutations in the C-terminal cytoplasmic region, which is not modelled in 7KR5, are indicated below the diagram. (B) Cytoplasmic Ca^2+^ concentration in HEK293 cells expressing mCherry-*Sm*Orai WT or the mutants H79S and V46A, in the absence of *Sm*STIM. (C) Cytoplasmic Ca^2+^ concentration in HEK293 cells coexpressing mCherry-*Sm*Orai WT or the indicated mutants and EGFP-*Sm*STIM. Ca²⁺ imaging experiments were independently repeated twice. The graphs show the mean of at least 20 cells from one representative experiment.

For mutations provisionally expected to activate the channel, we determined whether there was constitutive calcium influx when *Sm*ORAI was expressed without *Sm*STIM. Wildtype *Sm*ORAI was quiescent when expressed alone (Fig. 4B). In contrast, both the V46A channel and the H79S channel exhibited constitutive calcium influx (Fig. 4B). The corresponding replacements in *Hs*ORAI1 cause constitutive calcium influx, because loss of the V102 sidechain [*S. mansoni* V46] perturbs the positioning of the nonpolar segment in TM1 that serves as the channel gate, and loss of the H134 sidechain [*S. mansoni* H79] perturbs a TM1–TM2/3 interface that stabilizes the closed conformation of the pore (51, 52).

For expected blocking mutations, mutated *Sm*ORAI channels were coexpressed with *Sm*STIM, and the observed calcium influx was compared with the constitutive calcium influx carried by the wildtype *Sm*ORAI channel coexpressed with *Sm*STIM. *Sm*ORAI E50Q, F132A, V136N, and I189D/I192D mutations all caused a complete loss of *Sm*STIM-dependent calcium influx (Fig. 4C) The corresponding replacements in HsORAI1 impair channel function because the ring of E106 residues [E50] forms an essential calcium-binding site in the conductance pathway; the residues F187 [F132] and V191 [V136] have a role in the gating conformational change; and L273/L276 [I189/I192] are key residues in the C-terminal SOAR/CAD interaction site (51, 53, 54). There were minor differences in the findings for *Sm*ORAI and *Hs*ORAI1. For example, the pronounced but incomplete effect of the single I189D replacement was more reminiscent of the effect of replacements in the ORAI3 C-terminal site, where mutation of more than one nonpolar residue is needed to eliminate coupling to STIM1 (55). More pointedly, whereas the bulky nonpolar residue L194 in *Hs*ORAI1 at the extracellular end of the human TM1–TM2/3 hydrophobic cluster is essential for STIM1-evoked currents, the corresponding *S. mansoni* residue A139 is not conserved, indicating a difference between *Sm*ORAI and *Hs*ORAI1 in local structure in this region and in its contribution to channel gating.

We confirmed the findings by electrophysiological analysis of selected *Sm*ORAI mutants. We expressed *Sm*Orai alone in ORAI-TKO cells and found no constitutive or store depletion-activated current, reinforcing the conclusion that *Sm*Orai requires *Sm*STIM for its activity (Fig. 5 A,B). Like the corresponding human ORAI1 mutant (51, 52), *Sm*ORAI H79S carried a constitutive inward-rectifying current, with a reversal potential of +40 ± 6 mV (*n*=10) indicating a considerable degree of ion selectivity (Fig. 5 A,B). *Sm*Orai V46A exhibited a STIM-independent constitutive inward current at negative voltages and a large outward current— presumably carried by Cs^+^— at positive voltages, with a reversal potential of +19 ± 6 mV (*n*=10) (Fig. 5 A,B). This again mirrored the human counterpart (52, 56), and suggested a relatively nonselective cation channel. Coexpression of *Sm*Orai E50Q with *Sm*STIM gave no current, consistent with identification of E50 as part of the conserved selectivity filter of the schistosome CRAC channel (Fig. 5 C,D).

**Fig. 5.**
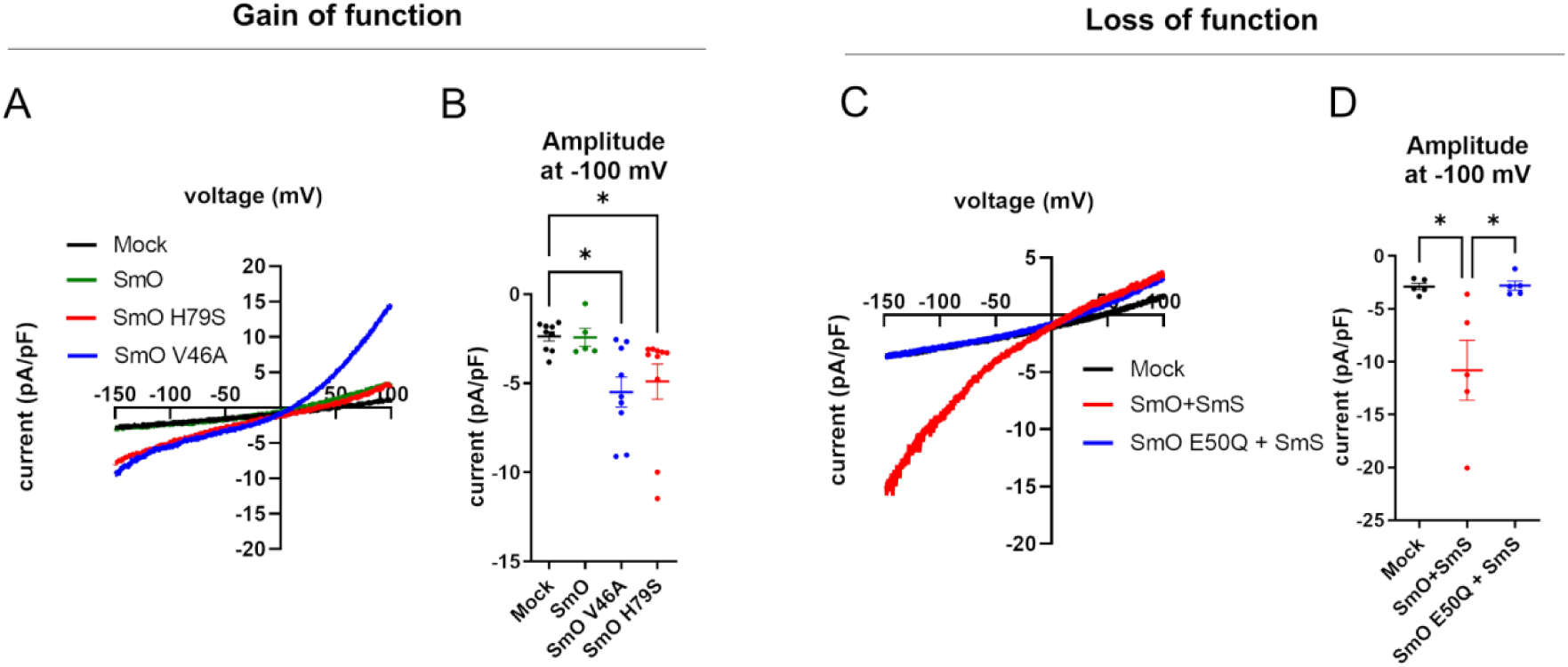
SmOrai H79S and V46A mutants generate smSTIM-independent constitutive currents. Average of I/V relationships (A) and current density at −100mV (B) from HEK-OraiTKO mock-transfected cells (N=9) and HEK-OraiTKO cells expressing either SmOrai (N=5), SmOrai H79S (N=10) or SmOrai V46A (N=9) mutants without smSTIM co-expression. Average of I/V relationships (C) and Current density at −100 mV (D) from HEK-OraiTKO mock-transfected cells (N=5) and HEK-OraiTKO cells co-expressing smSTIM with either smOrai (N=5) or smOrai E50Q (N=5) mutant.

In sum, this initial characterization demonstrated that *Sm*ORAI is a STIM-gated calcium channel, and indicated that basic features of the channel pore and the gating conformational change are conserved between *Sm*ORAI and *Hs*ORAI.

### *Sm*STIM is targeted to ER-plasma membrane junctions by its polybasic tail

A core function of STIM proteins is communication between the ER lumen and the plasma membrane. Human STIM1, when activated by a decrease in ER-luminal Ca^2+^ concentration, is targeted to ER-plasma membrane junctions by interactions of its C-terminal polybasic tail with the inner leaflet of the plasma membrane and by interactions of its SOAR/CAD domain with plasma membrane ORAI channels (57). The physical organization of *Sm*STIM resembles that of *Hs*STIM1— with a recognizable luminal EF-SAM domain, a single transmembrane segment, and a recognizable SOAR/CAD domain and polybasic segment in the cytoplasmic region (Fig. 6A; see also the *Sm*STIM-*Hs*STIM1 sequence alignment in Fig. S1) — making it plausible that similar mechanisms might target SmSTIM to ER-plasma membrane junctions.

**Fig. 6.**
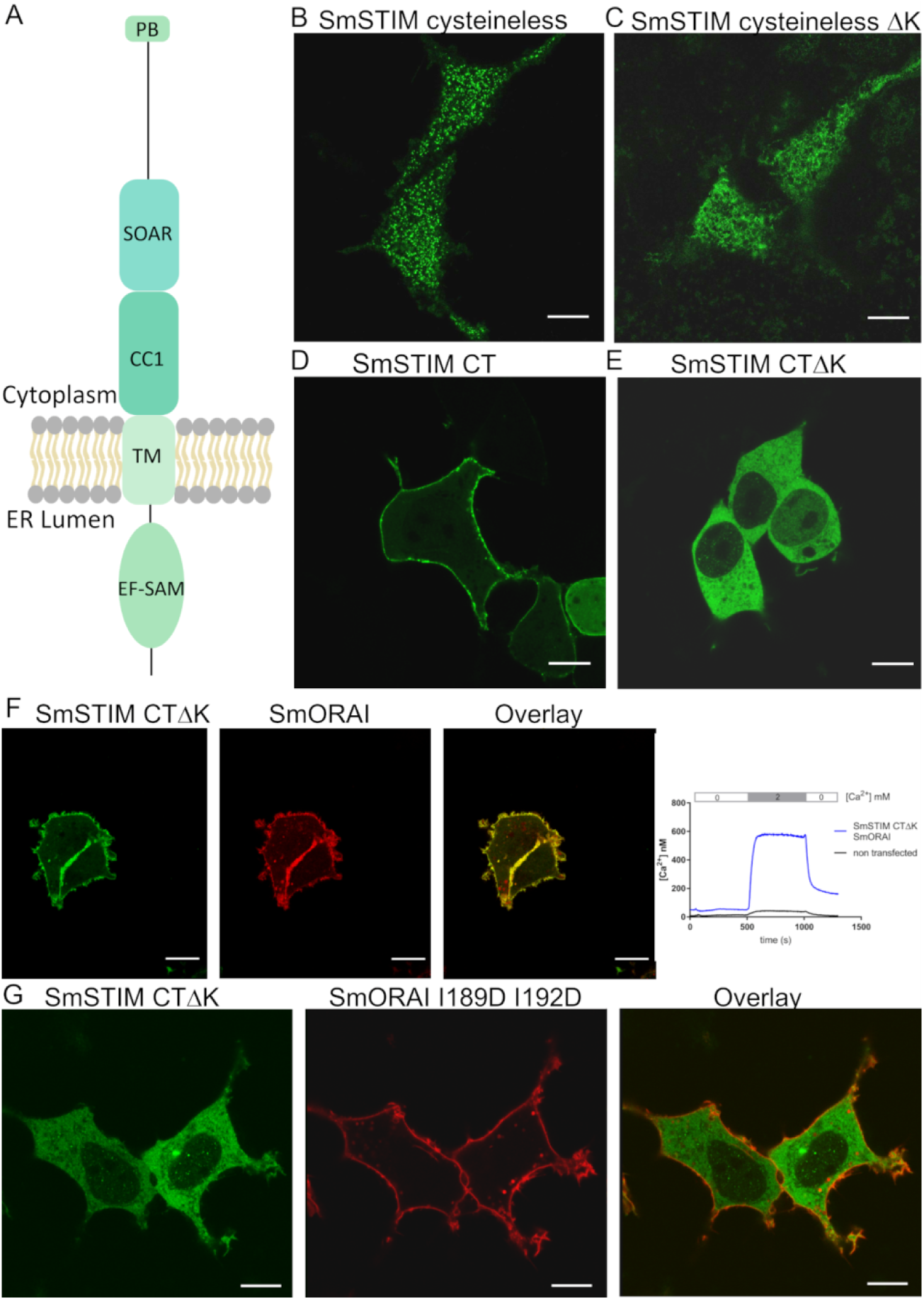
*Sm*STIM targeting to ER-PM junctions. (A) Schematic representation of *Sm*STIM depicting the luminal EFSAM domain (EF-SAM), single transmembrane helix (TM), SOAR/CAD domain, and C-terminal polybasic tail (PB). (B) Representative confocal image showing cysteineless EGFP-*Sm*STIM localized to junctions. (C) Representative confocal image showing cysteineless EGFP-*Sm*STIMΔK localized deeper in the ER. (D) EGFP-*Sm*STIM^CT^ including the entire *Sm*STIM cytoplasmic domain (residues 212-610) decorated the PM when expressed in the absence of *Sm*Orai. (E) EGFP-*Sm*STIM^CT^ΔK (residues 212-594) failed to decorate the plasma membrane when expressed in the absence of *Sm*Orai. (F) EGFP-*Sm*STIM^CT^ΔK coexpressed with mCherry-*Sm*Orai was recruited to the plasma membrane (*left*) and caused Ca^2+^ influx into the cells (*right*). (G) EGFP-*Sm*STIM^CT^ΔK coexpressed with mCherry-*Sm*Orai (I189D/I192D) did not decorate the plasma membrane. Scale bar, 10 µM. Experiments were independently repeated twice, with at least 10 cells imaged in each experiment. The images shown are representative.

Because wildtype *Sm*STIM was partially localized at ER-plasma membrane junctions in HEK293 cells, and partially localized deeper in the ER, it was difficult to quantitate its presence at junctions in order to study the mechanisms of targeting. We discovered that “cysteineless” *Sm*STIM (a construct in which all cysteines were replaced with serines except C200, a conserved residue in the transmembrane domain that was retained to minimize potential disruption of STIM structure) was reliably localized at junctions and was therefore easier to score for changes in localization. Cysteineless *Sm*STIM localized mainly to junctions even when expressed in the absence of *Sm*ORAI (Fig. 6B). Tellingly, omission of the 16-residue C-terminal polybasic segment from cysteineless *Sm*STIM—creating a protein truncated at residue 594, termed here cysteineless *Sm*STIMΔK— altered the cellular distribution of *Sm*STIM in the absence of *Sm*ORAI to a broad localization in the ER with no preference for junctions (Fig. 6C). This result indicates that the polybasic tail of *Sm*STIM, like that of *Hs*STIM1, is required for plasma membrane localization in the absence of ORAI.

In an independent approach, we assessed *Sm*STIM targeting to the plasma membrane by expressing the C-terminal cytoplasmic domain of *Sm*STIM— *Sm*STIM(212-610), termed *Sm*STIM^CT^ — as a soluble fragment untethered to the ER. Unlike *Hs*STIM1^CT^, which depends on ORAI for recruitment to the plasma membrane (57), the C-terminal fragment of wildtype *Sm*STIM decorated the plasma membrane even when expressed without *Sm*ORAI (Fig. 6D). ORAI-independent recruitment to the plasma membrane again depended on the STIM polybasic tail, since it was not observed with *Sm*STIM^CT^ ΔK [*Sm*STIM(212-594)] (Fig. 6E). Coexpression of *Sm*ORAI with *Sm*STIM^CT^ ΔK restored plasma membrane decoration by the STIM fragment (Fig. 6F, left panels) and led to constitutive calcium entry (Fig. 6F, right panel), indicating that additionally *Sm*STIM was targeted to the plasma membrane by a protein-protein interaction with *Sm*ORAI (see also Fig. S6). Consistent with the inability of *Sm*ORAI(I189D/I192D) to engage *Sm*STIM, as judged by calcium-entry [Figure 6C], introduction of those C-terminal replacements into *Sm*ORAI caused a loss of plasma membrane decoration by *Sm*STIM^CT^ ΔK (Fig. 6G), indicating that a primary interaction of *Sm*STIM occurs with the *Sm*ORAI C terminus.

In sum, wildtype *Sm*STIM expressed in HEK293 cells is partially localized to junctions, and cysteineless *Sm*STIM is largely localized to junctions. The current evidence does not cast light on whether *Sm*STIM typically localizes at junctions in schistosome cells, or whether *Sm*STIM activation is modulated differently in schistosome cells versus human cells. Nevertheless, analysis of cysteineless *Sm*STIM and *Sm*STIM fragments implies that *Sm*STIM can utilize the same two plasma-membrane targeting mechanisms as *Hs*STIM1.

### *Sm*STIM cytoplasmic domain favors constitutive activation

We obtained further insight into the constitutive activation of *Sm*STIM by focusing on the *Sm*STIM cytoplasmic domain. The *Sm*STIM cytoplasmic domain conferred resting localization partly at puncta on an *Hs*STIM1-*Sm*STIM^CT^ chimera (Fig. S7 A), and the isolated recombinant *Sm*STIM C-terminal domain was preferentially in an extended conformation, as assessed by the energy-transfer assay that previously established isolated recombinant *Hs*STIM1^CT^ is folded back on itself but can be extended by activation (58) (Fig. S7 B). Thus, as an isolated domain, *Sm*STIM^CT^ has more propensity toward activation than isolated *Hs*STIM1^CT^. The apparent preference of full-length *Sm*STIM for an activated conformation is, of course, a balance of diverse intramolecular and intermolecular interactions and cannot be referred solely to one STIM domain. However, as a practical matter, the data for this chimera suggested an approach that allowed us to dissect the calcium-sensing function of the *Sm*STIM luminal domain in the next section.

### *Sm*STIM ER-luminal domain senses ER calcium levels

A second core function of STIM proteins is ER-luminal calcium sensing, which in *Hs*STIM1 is coupled to a concerted conformational change extending from the luminal domain through the STIM cytoplasmic domain. Building on the findings of the previous section, we set out to examine calcium sensing by the *Sm*STIM ER-luminal domain independently of the dominant plasma membrane-targeting effect of the *Sm*STIM cytoplasmic domain, using a chimera of *Sm*STIM luminal domain and *Hs*STIM1^CT^ (Fig. 7A). This chimera exhibited localization throughout the ER in resting cells, relocalization to puncta upon store depletion, and store-dependent calcium entry (Fig. 7B). These observations establish that *Sm*STIM luminal domain is inherently capable of sensing a physiological decrease in calcium in the ER lumen and conveying this information to a STIM cytoplasmic domain.

**Fig. 7.**
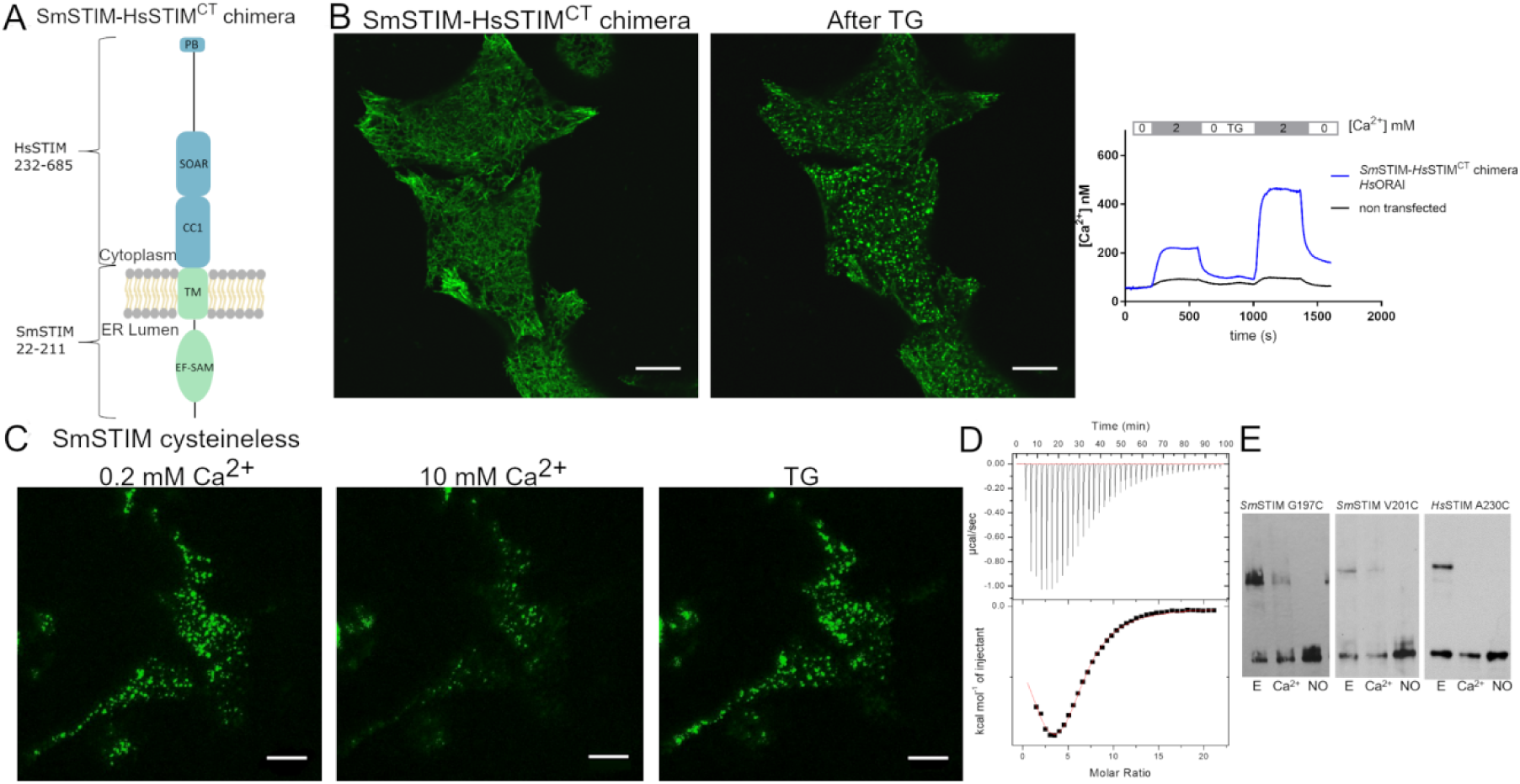
*Sm*STIM EFSAM domain is a calcium sensor. (A) Schematic representation of the chimera comprising the *Sm*STIM luminal domain and transmembrane segment fused to the *Hs*STIM1 cytoplasmic domain. (B) Wide distribution of the EGFP-*Sm*STIM-*Hs*STIM1^CT^ chimera across the ER in untreated cells (*left panel*) contrasts with preferential localization to ER-plasma membrane junctions after store depletion (*center panel*). The relocalization to junctions is associated with store-dependent calcium entry (*right panel*). (C) TIRF images of the same field of HEK293 cells expressing cysteineless EGFP-*Sm*STIM in a control extracellular containing 0.2 mM Ca^2+^ (*left panel*), after exchange to an extracellular solution containing 10 mM Ca^2+^ (*middle panel*), and then following stimulation with TG in nominally Ca^2+^-free solution (*right panel*). About 40% of HEK293 cells expressing cysteineless *Sm*STIM in the absence of *Sm*Orai showed the phenotype that exposure to an extracellular solution containing 10 mM Ca^2+^ reversed the junctional localization of cysteineless EGFP-*Sm*STIM. (D) Isothermal titration calorimetry of calcium binding to *Sm*STIM EFSAM-GrpE. *Upper panel*, Heat changes elicited by injections of 8 mM CaCl_2_ into a sample cell containing 80 µM protein. *Lower panel*, Integrated binding isotherm as a function of the Ca^2+^/protein molar ratio, after subtracting dilution heats. (E) Oxidative crosslinking by iodine of the single-cysteine mutants *Sm*STIM(G197C) and *Sm*STIM(V201C) in membranes isolated from HEK293 cells, in the presence of 0.5 mM EGTA (*E*) or 2 mM Ca^2+^ (*Ca^2+^*), monitored on a western blot as the upper dimer band. No crosslinking was observed in no-oxidation control samples (*NO*) where iodine was omitted. Crosslinking of *Hs*STIM(A230C) in the same experiment is shown for reference.

Interestingly in terms of calcium sensing, elevating extracellular calcium to 10 mM decreased junctional cysteineless *Sm*STIM in some cells (Fig. 7C). This would be most easily explained by ER calcium loading in those cells, shifting ER calcium concentration into the range where *Sm*STIM expressed in human cells undergoes a concerted conformational change to its inactive form, a sideline that we have not pursued further.

The protein-chemical correlates of calcium sensing have been defined in human STIM1 and STIM2 EF-SAM domains stabilized as soluble proteins by fusion with a *T. thermophilus* GrpE dimer (59). We applied the same methods to *Sm*EFSAM-GrpE. *Sm*STIM luminal domain calcium binding detected by isothermal titration calorimetry was biphasic in heat release, suggesting a conformational change coupled to calcium binding as in *Hs*EFSAM-GrpE; saturated in the range below 1 mM Ca^2+^; and corresponded to roughly 6 calcium-binding sites per monomer (Fig. 7D). A calcium-dependent conformational change of *Sm*EFSAM was further reflected in ANS binding and fluorescence as Ca^2+^ was titrated over the range 1 µM to 10 mM, and in visible changes in the sensitivity to digestion by limiting amounts of trypsin in the absence and presence of calcium (Fig. S8 A, B). The concerted conformational change in *Hs*STIM1 has been directly demonstrated in disulfide crosslinking experiments that probe the repositioning of TM helices of full-length STIM1 in ER membranes. Calcium sensing likewise altered *Sm*STIM TM-helix crosslinking in isolated HEK293 cell membranes, demonstrating an apposition of G197C and V201C residues directed by low Ca^2+^ (Fig. 7E). Note that schistosome STIM and human STIM1 have rather different TM sequences, and we have not confirmed that the detailed geometry of TM helix apposition is the same in the two cases.

These experiments, collectively, show that the calcium-sensing mechanism previously defined for human STIM1 is conserved in *Sm*STIM.

### *Sm*ORAI-*Hs*ORAI chimeras highlight differences in STIM-ORAI gating

Despite the shared mechanisms for calcium sensing, conformational change, and plasma-membrane targeting, and despite even the ability of *Hs*STIM1 to activate *Sm*ORAI, *Sm*STIM did not recognize *Hs*ORAI1. Thus, cysteineless *Sm*STIM localized to junctions in resting HEK293 cells, but coexpressed *Hs*ORAI1 did not colocalize with *Sm*STIM, nor did coexpression of *Hs*ORAI1 with *Sm*STIM increase calcium influx (Fig. 8A). Similarly, *Sm*STIM^CT^ ΔK failed to decorate the plasma membrane when coexpressed with *Hs*ORAI1, and failed to increase calcium influx, the negative outcome in each assay contrasting with the behavior of the same *Sm*STIM fragment when coexpressed with *Sm*ORAI (Fig. 8 B; compare Fig. 6F). The cross-interactions between human and schistosome STIM and ORAI proteins are summarized in (Fig. S9). These results indicate that it would be worth investigating further avenues to differential blockade of *Sm*STIM-*Sm*ORAI signalling, in addition to screening for selective pharmacological blockers of the *Sm*ORAI channel itself.

**Fig. 8.**
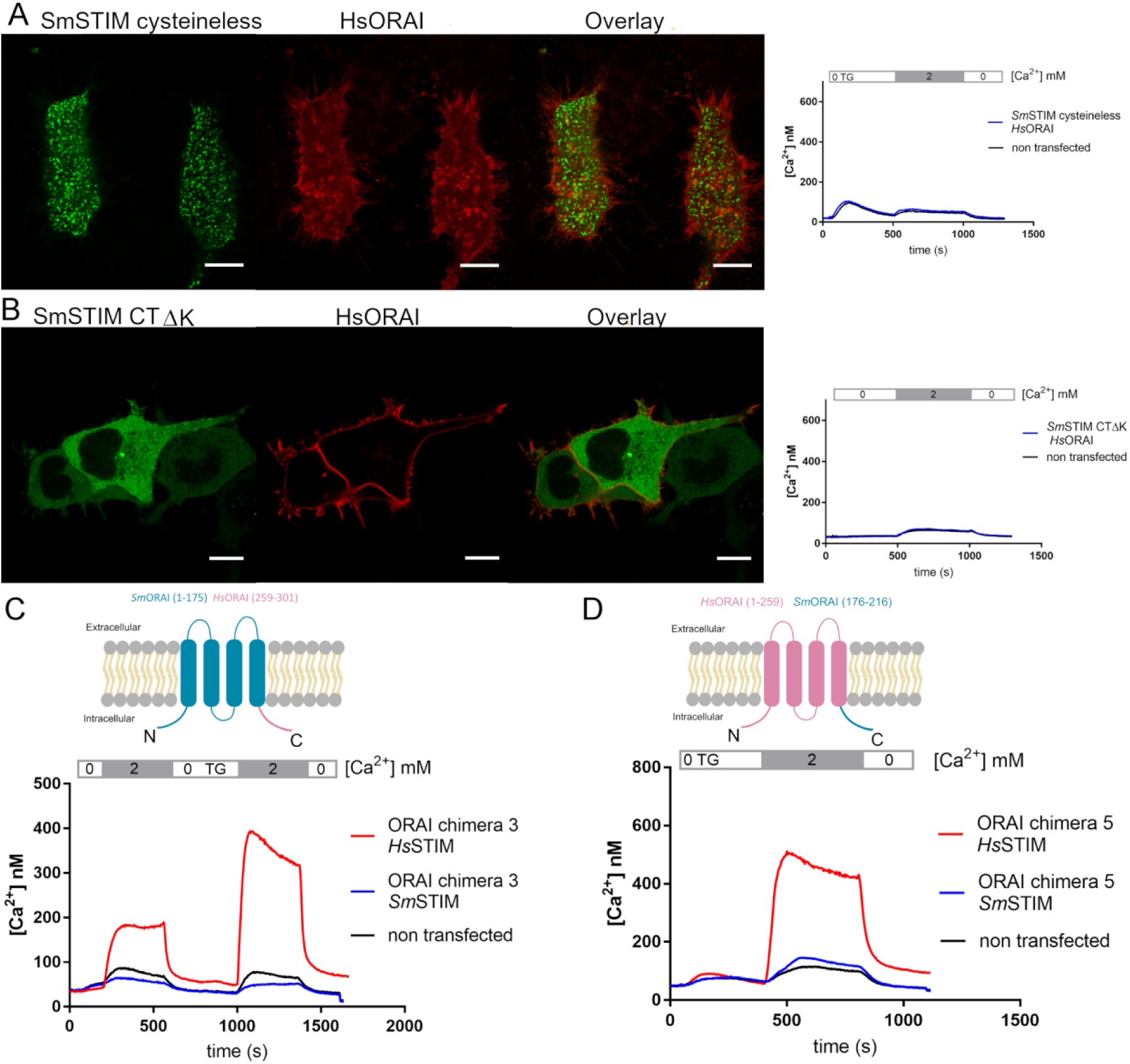
Insight into STIM-Orai gating from *Sm*Orai-*Hs*Orai chimeras. (A) *Sm*STIM does not recognize *Hs*Orai. Confocal images of representative HEK293 cells coexpressing cysteineless EGFP-*Sm*STIM and mCherry-*Hs*Orai (*left panels*) and cytoplasmic Ca^2+^ concentrations monitored in a separate sample of cells expressing the two proteins (*right panel*). Despite its preferential localization to ER-plasma membrane junctions, cysteineless EGFP-*Sm*STIM does not recruit mCherry-*Hs*Orai1 to junctions and does not trigger calcium entry. (B) EGFP-*Sm*STIM^CT^ΔK does not decorate the plasma membrane when coexpressed with mCherry-*Hs*Orai (*left panels*) and does not trigger Ca^2+^ influx into the cells through *Hs*Orai channels (*right panel*). Scale bars, 10 µM. (C) *Sm*STIM did not gate Orai chimera 3, either in resting cells or following store depletion. *Upper panel*, schematic representation of Orai chimera 3. *Lower panel*, Cytoplasmic calcium concentrations in cells coexpressing Orai chimera 3 and either *Hs*STIM1 (*red*) or *Sm*STIM (*blue*). The sequence of extracellular solutions applied is indicated above the graph of calcium concentrations. (D) *Sm*STIM did not gate Orai chimera 5. *Upper panel*, schematic representation of Orai chimera 5. *Lower panel*, Cytoplasmic calcium concentrations in cells coexpressing Orai chimera 5 and either *Hs*STIM1 (*red*) or *Sm*STIM (*blue*).

*Sm*ORAI-*Hs*ORAI chimeras implicated both the ORAI C terminus and a second region of the channel in gating. Perhaps predictably— in light of the STIM-ORAI decoupling effect of *Sm*ORAI C-terminal mutations (Fig. 4C and Fig. 6G) and the failure of *Sm*STIM to recognize *Hs*ORAI (Fig. 8A,B)— *Sm*STIM failed to gate ORAI chimera 3, which was derived from *Sm*ORAI by replacing its C terminus with *Hs*ORAI1^CT^ (Fig. 8C). The chimeric channel was functional, since it could be gated by *Hs*STIM1. In particular, the latter experiment established that the human ORAI1 C terminus could engage productively in a gating conformational change with the remainder of a channel composed of the schistosome sequence. The deficit in *Sm*STIM–ORAI chimera 3 signalling can be explained by a failure of *Sm*STIM to recognize the HsORAI1 C terminus. The overall deficit in *Sm*STIM–*Hs*ORAI1 functional communication was not limited to the ORAI C terminus, though, because *Sm*STIM also failed to gate the converse ORAI chimera 5, in which the C terminus of *Hs*ORAI1 had been replaced by *Sm*ORAI1^CT^ (Fig. 8D). Again, the schistosome ORAI C terminus was able to engage productively with the remainder of a channel composed of the human sequence and the chimeric channel was functional, since *Hs*STIM1 gated ORAI chimera 5. These results indicate that *Sm*STIM must recognize a second region of the ORAI channel in order to effect gating.

We asked whether the ORAI N terminus affected gating by the two species of STIM differentially, comparing activation of wildtype *Sm*ORAI and ORAI chimera 1 (*Hs*ORAI1^NT^-*Sm*ORAI), either by *Sm*STIM or by *Hs*STIM1. Strikingly, engrafting the human ORAI1 N terminus in place of the schistosome ORAI N terminus resulted in a marked decrease in activation of the channels by *Sm*STIM, but a marked increase in activation by *Hs*STIM1 (Fig. 9A). The data are not consistent with the hypothesis that ORAI channel gating depends only on STIM interaction with the ORAI C terminus (60). To rule out the possibility that the more extended N terminus of *Hs*ORAI1 interferes selectively with productive *Sm*STIM interaction with the chimeric channels, we repeated the experiment by engrafting only the portion of the human ORAI1 N terminus beginning at residue M64, an alternative translation start site of *Hs*ORAI1 (61, 62). In this case, the engrafted portion of human ORAI1, *Hs*ORAI1(64–80), is shorter than the native N-terminal sequence of *Sm*ORAI (Fig. 9B). The truncation did not improve the ability of *Sm*STIM to activate the chimeric channels (Fig. 9B). The most straightforward interpretation of the differential activation of the N-terminal *Sm*ORAI chimera by *Sm*STIM and *Hs*STIM1 would be that the STIM cytoplasmic domain interacts with the ORAI N-terminal segment— a possibility that has been debated in the literature (47, 53, 57, 60, 63–69) — but there are alternative interpretations and definitive examination of this possibility would have been a digression from our goals here. The conclusion relevant to our interest in pharmacological targeting of STIM-ORAI signalling in parasitic worms is that the physiological gating interactions of *Sm*STIM and *Hs*STIM1 with their cognate ORAI channels differ in detail, indicating differences in structure that could enable selective pharmacological blockade of the schistosome channel.

**Fig. 9.**
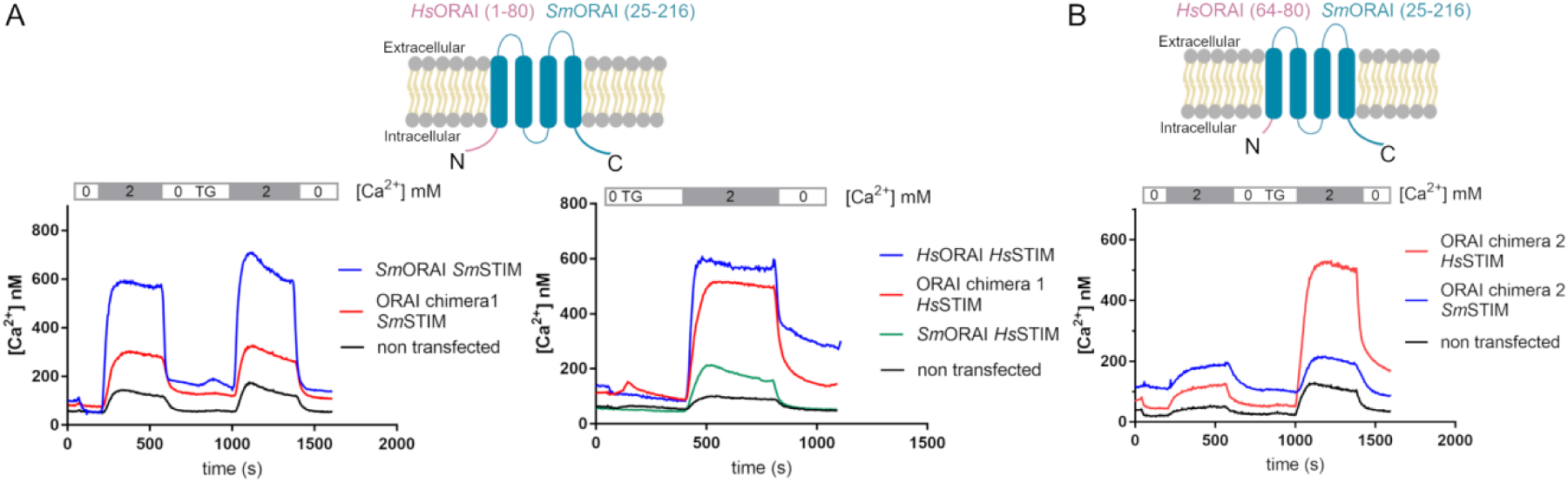
The Orai N terminus affects channel gating. (A) *Upper panel*, schematic representation of Orai chimera 1. *Lower left panel*, Cytoplasmic calcium concentrations in cells coexpressing *Sm*STIM and either *Sm*Orai (*blue*) or Orai chimera 1 (*red*). *Lower right panel*, Cytoplasmic calcium concentrations in cells coexpressing *Hs*STIM1 with *Hs*Orai1 (*blue*), Orai chimera 1 (*red*), or *Sm*Orai (*green*). (B) Schematic representation of Orai chimera 2, consisting of *Hs*Orai1(64–80) fused to *Sm*Orai(25-216). Cytoplasmic calcium concentrations measured in HEK293 cells coexpressing mCherry-Orai chimera 2 and either *Hs*STIM (*red*) or *Sm*STIM (*blue*). The sequence of extracellular solutions applied is indicated above the graph of calcium concentrations.

### *Sm*STIM-*Sm*ORAI exhibits distinctive sensitivity to pharmacological blockers

To test directly whether pharmacological blockers can discriminate between the schistosome and human ORAI channels, we examined the effects of four selective inhibitors of the human CRAC channel developed by CalciMedica (70) and the effects of the pyrazole inhibitor BTP2 (71, 72) on HEK293 cells coexpressing either *Hs*STIM1 and *Hs*ORAI1 or *Sm*STIM and *Sm*ORAI. Either CM2748 or CM4308 at 1 µM substantially blocked thapsigargin-evoked calcium influx through both the human and schistosome ORAI channels, compared to the influx observed in parallel DMSO vehicle-treated controls (Fig. 10A and 10B). In contrast, CM6325 at 1 µM fully blocked calcium influx into cells expressing *Hs*STIM1 and *Hs*ORAI1, but had little effect against cells expressing *Sm*STIM and *Sm*ORAI (Fig. 10C). CM5480 also blocked only the human channel. Titration established that CM6325 fully blocked calcium influx through the human ORAI1 channel at 200 nM, but produced only a partial block of the schistosome ORAI channel at 10 µM (Fig. 10D). BTP2 exhibited a similar large difference in efficacy (Fig. 10E).

**Fig. 10.**
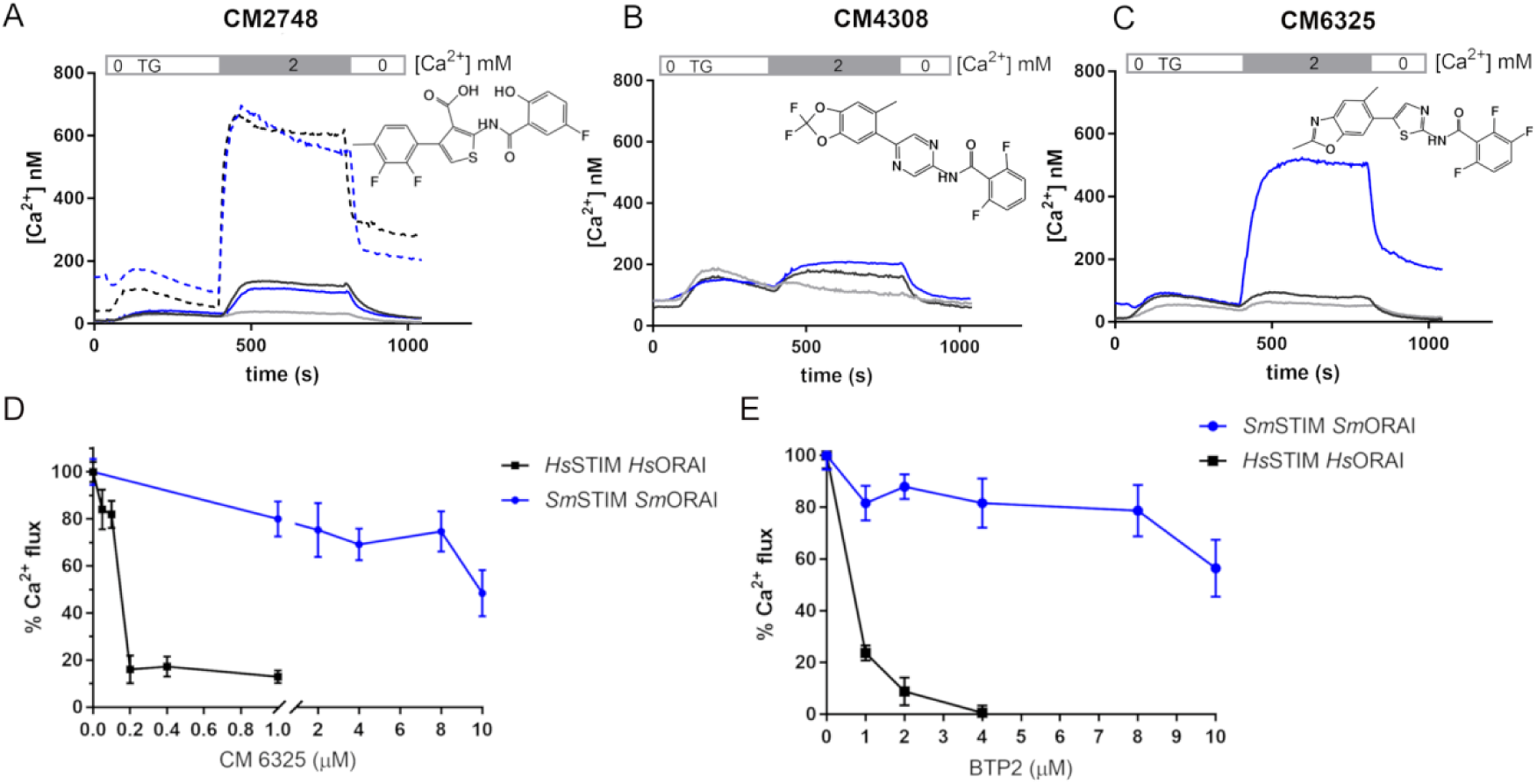
*H. sapiens* and *S. mansoni* CRAC channels display pharmacological differences. (A) Cytoplasmic calcium concentrations measured in HEK293 cells coexpressing EGFP-*Hs*STIM1 and mCherry-*Hs*Orai1 (*black curves*) or coexpressing EGFP-*Sm*STIM and mCherry-*Sm*Orai (*blue curves*). The dashed curves correspond to control cells preincubated with DMSO vehicle. The cells had been preincubated with 1 µM CM2748 for 1h. The sequence of extracellular solutions applied is indicated above the graph of calcium concentrations. CM2748 was present during the calcium measurements. Data for non-transfected cells in the same cultures, defined by the absence of detectable GFP fluorescence are plotted as a *gray solid curve*. (B–C) Cytoplasmic calcium concentrations measured as described in (A), but in the presence of the CalciMedica compounds indicated at the top in each panel. Measurements shown in panels (A)–(C) were performed on the same day, hence the data shown in panel (A) for control cells preincubated with DMSO vehicle serve as a reference for all four panels. (D–E) Titration examining the effect of CM6325 or BTP2, as indicated, on Ca^2+^ entry. HEK293 cells coexpressing *Hs*STIM1 and *Hs*Orai1 (*black*) or *Sm*STIM and *Sm*Orai (*blue*) were preincubated with CM6325 or BTP2 at the indicated concentrations for 1h, and the measurements were performed in the continuing presence of the inhibitor. Percentages plotted are relative to the signal from cells expressing the same pair of proteins— schistosome or human— and preincubated with DMSO. Error bars represent the standard error of the mean (SEM) from at least 20 cells.

While it is not surprising that compounds originally selected as inhibitors of the human ORAI1 channel display a bias toward blocking *Hs*ORAI1 over *Sm*ORAI, these data further document that there are pharmacological differences between the schistosome and human ORAI channels, which might be exploited to identify schistosome-selective ORAI channel blockers.

## Discussion

We describe here the first characterization of a parasitic helminth CRAC channel, from *Schistosoma mansoni*. Its underlying mechanisms resemble those of other STIM-ORAI pairs in STIM calcium sensing, STIM targeting to ER-plasma membrane junctions, STIM-ORAI interaction, ORAI pore architecture, and ORAI gating mechanism.

In the absence of a workable schistosome cell preparation, we cannot verify that STIM-ORAI signalling is silent in resting schistosome cells. However, the fact that the calcium-sensing function of the ER-luminal domain is retained in *Sm*STIM suggests that ER calcium controls an inactive–active switch in schistosome cells, as it does in mammalian cells. In this case, the propensity of the isolated recombinant C-terminal of *Sm*STIM to assume an extended conformation would have to be countered in resting schistosome cells by constraints such as STIM-protein interactions or STIM-lipid interactions that keep *Sm*STIM in its inactive state, or by a relative shortfall of factors that favor activation. Our data indicate that *Sm*STIM is balanced just beyond the edge of the inactive–active transition even when it is expressed in mammalian cells, since we have shown that the constitutive near-plasma membrane localization of *Sm*STIM in mammalian cells is ‘curable’ in many cells simply by elevating the extracellular calcium concentration.

The behavior of *Sm*STIM in our experiments resembles, at first glance, that of *C. elegans* STIM (*Ce*STIM), which also localizes constitutively in near-plasma membrane puncta when expressed in HEK293 cells (73, 74). However, unlike the schistosome protein, *Ce*STIM is not fully activated under these conditions, since it recruits ORAI to puncta and triggers calcium influx only after ER calcium store depletion (73–75). There are probably no deep mechanistic lessons to be learned by comparing the two cases, because *S. mansoni* (a flatworm) and *C. elegans* (a roundworm) are very distant evolutionarily, and because *Sm*STIM has no more sequence similarity with *Ce*STIM than it has with *Hs*STIM1.

The finding that replacing the N-terminal region of *Sm*ORAI with the human ORAI N-terminal peptide has opposite effects on channel activation by *Sm*STIM and *Hs*STIM1 refocuses attention on the possibility that STIM engages the ORAI N terminus directly at some time during channel activation. There is ample evidence that the ORAI N terminus in isolation binds STIM (47, 57, 63, 64, 69), but whether such an interaction participates in gating the intact ORAI channel is not settled (76–79). Our observations do not settle the issue, but they do rule out the very simplest model, that binding of STIM to the ORAI C terminus is by itself sufficient for full channel gating (60).

Our central findings are that *S. mansoni* and *H. sapiens* STIM and ORAI have close functional similarities, but that there are nevertheless promising pharmacological distinctions between the schistosome and human CRAC channels. We tested four compounds developed as selective blockers of human CRAC channels on HEK293 cells expressing schistosome or human CRAC channels. Importantly, only two of the four compounds (CM2748 and CM4308) were efficient in inhibiting calcium influx through both schistosome and human CRAC channels, whereas the two others (CM5480 and CM6325) had very low efficacy against *Sm*CRAC channel-expressing cells, even though they were fully efficacious against *Hs*CRAC channel-expressing cells. This differential sensitivity to inhibitors indicates that it would be worthwhile to screen for selective *S. mansoni* CRAC channel inhibitors, and suggests that *Sm*STIM and *Sm*ORAI are potentially new targets for the prevention and treatment of schistosomiasis.

## Materials and Methods

### Plasmids

Two splice forms of *Sm*Orai were found in the genome of *S. mansoni*, all the experiments described in this paper were performed with the longer isoform, Smp_076650.2. Two partial sequences were found for *Sm*STIM (Smp_174640 e Smp_180230) and the full length protein was obtained through search in the EST (expressed sequence tags) database. RNA extracted from *S. mansoni* adult worms was used to synthesize cDNA, which was then used as template in PCR reactions to amplify *Sm*STIM and *Sm*Orai.

The full length sequences were cloned into pGEM-T (Promega) propagation vector and subcloned into mammalian or bacterial expression vectors. For mammalian cell expression, *Sm*STIM was subcloned into pCMV6-XL5 with EGFP tagged at the N terminus. *Sm*Orai was subcloned in the MO91 vector with mCherry fusion protein at the N terminus.

STIM and Orai chimeras were constructed by PCR amplifying the domains from *H. sapiens* and *S. mansoni* separately and assembling them using the NEBuilder^®^ HiFi DNA Assembly Cloning Kit (New England Biolabs). Orai chimeras were in the MO91 vector and STIM chimeras in the pCMV6-XL5 vector. Mutations were introduced into mCherrry-SmOrai or HA-*Sm*STIM with QuikChange site directed mutagenesis kit (Agilent).

*Hs*STIM and *Hs*Orai were also in the plasmids pCMV6-XL5 and MO91, respectively. The cytoplasmic constructs of N-terminally tagged EGFP-*Sm*STIM (EGFP-*Sm*STIM^CT^ and EGFP-*Sm*STIM^CT^ ΔK were subcloned into pEGFP-C2. The crosslinking experiments were performed with HA-SmSTIM mutants subcloned into pcDNA3.1. The *Sm*STIM constructs for bacteria expression were subcloned in pet28a (*Sm*STIM EF-SAM-GrpE) or pProEX (GFP-*Sm*STIM-LBT) vectors, in the latter, a lanthanide-binding tag (GGFIDTNNDGWIEGDELLLEEG) was added to the C terminus of the protein.

### Confocal and TIRF microscopy

HEK293 cells were transfected with EGFP-*Sm*STIM and mCherry-*Sm*Orai or the specified construct using lipoafectamine 2000 (Thermo Fisher) according to the manufacturer*’s* instructions. After transfection, the cells were transferred to DMEM with low calcium (0.2 mM), supplementented with 10% FBS and antibiotics (Pen-Strep) and 10 µM LaCl_3_, added to the media to protect the cells from calcium overload. Cells were placed in Poly-D lysine (Sigma) pre-coated coverslips or on 35 mm glass-bottom dishes (MatTek) and imaged 24 h post transfection. Confocal images were acquired on a Zeiss LSM 880 microscope and TIRF images on a Nikon Ti-E microscope equipped with a PerfectFocus (PFS) module for TIRF imaging. The non-transfected cells were present in the same culture as transfected cells, but showed no detectable GFP fluorescence. Confocal and TIFR images presented in Figures 1, 6, 7, 8, S6, and S7 are representative of at least two independent experiments, in which a minimum of 10 cells were imaged in each experiment.

### Ca^2+^ influx measurements

Single cell intracellular Ca^2+^ levels were measured in HEK293 cells transiently expressing EGFP-*Sm*STIM, mCherry-*Sm*Orai or variants described in each experiment. The transfected cells on poly-D-lisine coated coverslips (18 mm) were loaded with Fura2 AM (Thermo Fisher catalog number F1221) in Ringer buffer (155 mM NaCl, 4.5 mM KCl, 10 mM glucose, 5 mM Hepes pH 7.4, 3 mM MgCl_2_) and 0.002% Pluronic F-127 (0.2 mm Ca^2+^were added in Ringer’s buffer for constitutive active constructs and 2 mM otherwise) and incubated for 1 h at 37 °C under 5% CO_2._ Cells were then washed twice in Ringer’s buffer and placed into a chamber to be imaged in an Olympus IX 71 microscope with the Polychrome V monochromator (TILL Photonics). Cells were alternately illuminated at 340 nm and 380 nm every 4 s and perfused with buffers containing no calcium, 1µM thapsigargin or 2 mM Ca^2+^, as indicated in each experiment. The average ratios obtained from at least 20 transfected cells were converted into Ca^2+^ concentration according to:

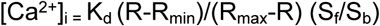

Where K_d_ = 220 nM, R_min_ and R_max_ were obtained from in situ calibration of HEK293 cells loaded with Fura2 (3 µM) and S_f_ and S_b_ are the emission intensities at 380 nm for the Ca^2+^ free and bound Fura2, respectively.

For calibration, cells were treated with 10 µM ionomycin in Ringer buffer supplemented with either 10 mM EGTA to obtain R_min_ (Ca²⁺-free condition) or 20 mM CaCl₂ to obtain R_max_ (Ca²⁺-saturated condition). Data analysis was performed using TillVision software (TILL Photonics). Experiments were repeated at least twice.

### Whole-cell patch clamp recordings

CRAC currents were measured using the whole cell configuration of the patch clamp technique in HEK-OraiTKO cells expressing various combinations of human and Schistosoma STIM and Orai as previously described (Zhang et al. 2019). Patch pipettes (2-4 MΩ) were pulled from borosilicate glass capillaries (World Precision Instruments, USA) with P-1000 Flamming/Brown micropipette puller (Sutter Instrument Company, USA). Data acquisition was performed using the Axopatch 200B, the Digidata 1440A (Molecular Devices, USA) and was monitored with pCLAMP10 software. Before recordings, cells transfected with Schistosoma STIM/Orai were trypsinized and incubated for 2 hours in low-Ca^2+^ media in round coverslips pre-treated with poly-L-Lysine to allow cell adhesion. Then, media was washed with the bath solution containing Na-methanesulfonate (115 mM), CsCl (10 mM), MgSO4 (1.2 mM), CaCl2 (20 mM), D-Glucose (10 mM), HEPES (10 mM). We adjusted pH at 7.4 with NaOH. We used Divalent-free solution (DVF), containing Na-Metanesulfonate (135 mM), HEDTA (10 mM), EDTA (1 mM), HEPES (10 mM), and with adjusted pH at 7.4 using NaOH, to measure Na^+^-mediated-SmOrai/SmSTIM currents. Pipettes were filled with a solution containing Cs-Methanesulfonate (115 mM), Cs-BAPTA (20 mM), MgCl2 (8 mM), HEPES (10 mM). We adjusted pH at 7.2 with CsOH. We included 8 mM MgCl_2_ in the pipette solution to inhibit TRPM7 currents. To measure Calcium Dependent Inactivation (CDI), we used 10 mM EGTA in the pipette instead of 20 mM Cs-BAPTA. We selected cells with tight seals (>3 GΩ) and with <15 MΩ series resistance for recording. For the ramp protocol, cells were maintained at +30 mV during the recordings and were subjected to reverse voltage ramps ranging from +150 mV to −150 mV, lasting 250 ms every 3s. For IV steps, cells were maintained at 0 mV and steps of +20 mV were applied from −140 mV to +140 mV, lasting 250 ms. We used acute administration of Gd^3+^ (concentration range from 1 µM to 50 µM) to inhibit SmOrai/SmSTIM-mediated currents. Currents in response to multiple ramps were collected and data analysis was performed by picking one single time point for each cell, which represent a stable signal that was chosen around 200-300s after break-in.

### CalciMedica Compounds

Stock solutions of CaMedica compounds CM2748, CM4308, CM6325 and CM5480 were freshly prepared in DMSO at 10 mM. Cells co-expressing either EGFP-*Sm*STIM and mCherry-*Sm*Orai or EGFP-*Hs*STIM and mCherry-*Hs*Orai were incubated with 1 µM of each compound (or the specified concentration for each experiment) for at least 1h and submitted to Ca^2+^ influx measurements. The incubation of the cells with Fura2, washing and perfusion were all performed in the presence of the specified compound. Each experiment was independently repeated at least three times, with a minimum of 20 cells analyzed per condition.

### Crosslinking experiments

Crosslinking expreriments were carried out according to our previus procedures (Hirve et al., 2018). Briefly, *Sm*STIM single cysteineless mutants were subcloned into pcDNA 3.1 vector fused to 3x HA tag at the N terminus. HEK293 cells were transfected with *Sm*STIM or *Hs*STIM constructs using Lipofectamine 2000. Cells were kept under 5% CO2 in DMEM supplementented with 10% FBS and antibiotics (Pen-Strep) and harvested 24 h after transfection. The cell pellet was suspended in 25 mM Tris pH 7.5, 25 mM NaCl, DNase I (12.5U/mL), 0.3 mM DTT and protease inhibitors. EGTA (0.5 mM) or Ca^2+^ (2 mM) was added to the buffer and the cells lysed by passage through a 25G syringe needle. The cell membranes were then collected after spinning at 100,000 g in a Beckman airfuge for 20 min, suspended in buffer (25 mM Tris pH 7.5, 150 mM NaCl, 0.3 mM DTT) containing EGTA or Ca^2+^ and oxidized with iodine (prepared by 10 times dilution of Lugol solution) for 10 min at 4 °C. The reaction was quenched by iodoacetamine (100 mM in 60 mM Tris pH 6.8 and 5x nonreducing loading buffer). Samples were then heated for 10 min at 55°C and analyzed by SDS-PAGE in precast 3-8% NuPAGE Bis-Tris gels. Western blots were carried out incubating with mouse anti-HA antibody (Sigma-Aldrich; catalog number H3663) (1:3000 dilution) and secondary HRP goat anti-mouse (1:4000 dilution) antibody (Sigma-Aldrich). ECL substrate (Perkin Elmer) was used to develop the blots.

### Protein expression and purification

*Sm*STIM EF-SAM-GrpE in pET28 (containing a His tag at the N terminus) was transformed into *E. coli* BL21 strain. Cells were grown at 37°C and 220 RPM shaking and the protein expression induced with 1 mM IPTG (isopropyl-β-D-thiogalactopyranoside) when the OD_600_ of the culture reached 0.6. The expression was carried out a 16 °C for 12-16h.

Bacterial cells were harvested and the pellet suspended into 30 mL of buffer 50 mM Tris pH 7.5, 150 mM NaCl, 5% Glycerol, 3 mM β-mercaptoethanol, protease inhibitors (Roche) and 20 mM CaCl_2._ Cells were lysed by sonication and cell debris separated by centrifugation (15,000 RPM 30 min). The supernatant was applied to Ni-NTA resin (Qiagen). Elution was carried out with the same buffer containing 300 mM imidazole, after washing steps with buffer containing 40 mM imidazole. A second purification step followed on a gel filtration Superdex 200 Increase 10/300 GL column (GE Healthcare) and proteins were eluted in lysis buffer containing 20 mM Ca^2+^ or in Ca^2+^ free buffer treated with Chelex-100 resin (Bio-Rad), in this case the protein was applied twice into the column to remove the calcium.

### Isothermal Titration Calorimetry

For ITC measurements, the *Sm*STIM EF-SAM-GrpE protein (80 µM) was placed in the sample cell of a MicroCal ITC-200 microcalorimeter (Malvern) and titrated at 20 °C with 8 mM CaCl_2,_ 1 µL aliquots of the ligand in the syringe was added to the sample cell until saturation was observed (the heat release corresponding only to CaCl_2_ dilution into buffer). Both protein and ligand were diluted in the same buffer (50 mM Tris pH 7.5, 150 mM NaCl, 5% Glycerol) and the dilution heat of the CaCl_2_ was subtracted for data analysis, performed in the Origin7 software (MicroCal). ITC measurements were performed in two independent experiments.

### Trypsin digestion and ANS experiments

Trypsin diluted at 5 µg/mL (Hamptom Research) was added to purified *Sm*STIM EF-SAM-GrpE protein and incubated for 1h at 37 °C. Loading buffer was added to the samples, heated at 90 °C for 10 min and analyzed into SDS-PAGE (4–12% NuPAGE gel; Life Technologies)

The 8-anilino-1-naphthalenesulfonic acid (ANS) assay was performed with 0.5 µM of *Sm*STIM EF-SAM-GrpE protein in chelex-treated buffer (50 mM Tris pH 7.5, 150 mM NaCl, 5% Glycerol) and 20 µM of ANS. The ANS probe was excited at 350 mm and the fluorescence emission spectra collected from 400-600 nm using a QuantaMaster 40 spectrofluorometer (Photon Technology International). Increasing amounts of calcium were added to the buffer and the spectra collected after 10 min incubation.

### LRET assay

LRET assay was performed as described by Zhou et al 2013. Briefly, GFP-*Sm*STIM (212-489)-LBT or GFP-*Hs*STIM (273-685)-LBT were expressed and purified as previously described for *Sm*STIM EF-SAM-GrpE, but without CaCl_2_ added to the buffer. The measurements were performed at 4°C in 80% glycerol buffer using QuantaMaster 40 spectrofluorometer (PTI). Tb^3+^ was excited at 280 nm and the luminescence or Tb^3+^-sensitized acceptor emission monitored from 400-600 nm.

## Acknowledgments

We would like to thank the late Prof. Ricardo DeMarco, who has left us too early, for his invaluable contribution at the beginning of this work. We acknowledge Dr Anjana Rao for her comments on the manuscript. This work was funded by National Institutes of Health grants AI084167, AI040127, AI109842 to PGH and NIH Grant R35HL150778 to MT. AEZ was supported in part by postdoctoral fellowship 2016/12505–8 from the São Paulo Research Foundation (FAPESP).

## References

1. P. J. Hotez, et al., Global Burden Disease Study 2010: Interpretation Implications Neglected Tropical Diseases. PLoS Neglected Tropical Diseases 8 (2014).

2. F. R. Martins-Melo, et al., The burden of Neglected Tropical Diseases in Brazil, 1990-2016: A subnational analysis from the Global Burden of Disease Study 2016. PLoS Negl. Trop. Dis. 12, e0006559 (2018).

3. D. P. McManus, et al., Schistosomiasis. Nat. Rev. Dis. Primers 4, 13 (2018).

4. C. M. Gower, L. Vince, J. P. Webster, Should we be treating animal schistosomiasis in Africa? The need for a One Health economic evaluation of schistosomiasis control in people and their livestock. Trans. R. Soc. Trop. Med. Hyg. 111, 244–247 (2017).

5. P. Adeyemo, et al., Estimating the financial impact of livestock schistosomiasis on traditional subsistence and transhumance farmers keeping cattle, sheep and goats in northern Senegal. Parasit. Vectors 15, 101 (2022).

6. E. Léger, et al., Prevalence and distribution of schistosomiasis in human, livestock, and snail populations in northern Senegal: a One Health epidemiological study of a multi-host system. *Lancet Planet*. Health 4, e330–e342 (2020).

7. O. P. Aula, D. P. McManus, M. K. Jones, C. A. Gordon, Schistosomiasis with a focus on Africa. Trop. Med. Infect. Dis. 6, 109 (2021).

8. N. C. Lo, et al., Review of 2022 WHO guidelines on the control and elimination of schistosomiasis. Lancet Infect. Dis. 22, e327–e335 (2022).

9. M. Debesai, M. Russom, Praziquantel and risk of visual disorders: Case series assessment. PLoS Negl. Trop. Dis. 14, e0008198 (2020).

10. P. T. Hoekstra, et al., Limited efficacy of repeated praziquantel treatment in Schistosoma mansoni infections as revealed by highly accurate diagnostics, PCR and UCP-LF CAA (RePST trial). PLoS Negl. Trop. Dis. 16, e0011008 (2022).

11. L. Pica-Mattoccia, D. Cioli, Sex- and stage-related sensitivity of Schistosoma mansoni to in vivo and in vitro praziquantel treatment. Int. J. Parasitol. 34, 527–533 (2004).

12. A. A. Sabah, C. Fletcher, G. Webbe, M. J. Doenhoff, Schistosoma mansoni: chemotherapy of infections of different ages. Exp. Parasitol. 61, 294–303 (1986).

13. N. Vale, et al., Praziquantel for schistosomiasis: Single-drug metabolism revisited, mode of action, and resistance. Antimicrob. Agents Chemother. 61 (2017).

14. R. Bergquist, J. Utzinger, J. Keiser, Controlling schistosomiasis with praziquantel: How much longer without a viable alternative? Infect. Dis. Poverty 6, 74 (2017).

15. J. T. Moreira-Filho, et al., Schistosomiasis drug discovery in the era of automation and artificial intelligence. Front. Immunol. 12, 642383 (2021).

16. M. Berriman, et al., The genome of the blood fluke Schistosoma mansoni. Nature 460, 352–358 (2009).

17. Schistosoma japonicum Genome Sequencing and Functional Analysis Consortium, The Schistosoma japonicum genome reveals features of host-parasite interplay. Nature 460, 345–351 (2009).

18. F. Luo, et al., An improved genome assembly of the fluke Schistosoma japonicum. PLoS Negl. Trop. Dis. 13, e0007612 (2019).

19. A. V. Protasio, et al., A systematically improved high quality genome and transcriptome of the human blood fluke Schistosoma mansoni. PLoS Negl. Trop. Dis. 6, e1455 (2012).

20. A. J. Stroehlein, et al., High-quality Schistosoma haematobium genome achieved by single-molecule and long-range sequencing. Gigascience 8 (2019).

21. N. D. Young, et al., Whole-genome sequence of Schistosoma haematobium. Nat. Genet. 44, 221–225 (2012).

22. G. Sankaranarayanan, M. Berriman, G. Rinaldi, An uneven race: genome editing for parasitic worms. Nat. Rev. Microbiol. 19, 621 (2021).

23. W. Ittiprasert, et al., Programmed genome editing of the omega-1 ribonuclease of the blood fluke, Schistosoma mansoni. Elife 8 (2019).

24. G. Sankaranarayanan, et al., Large CRISPR-Cas-induced deletions in the oxamniquine resistance locus of the human parasite Schistosoma mansoni. Wellcome Open Res. 5, 178 (2021).

25. H. You, et al., CRISPR/Cas9-mediated genome editing of Schistosoma mansoni acetylcholinesterase. FASEB J. 35, e21205 (2021).

26. G. Wendt, et al., A single-cell RNA-seq atlas of Schistosoma mansoni identifies a key regulator of blood feeding. Science 369, 1644–1649 (2020).

27. J. Wang, et al., Large-scale RNAi screening uncovers therapeutic targets in the parasite Schistosoma mansoni. Science 369, 1649–1653 (2020).

28. S. R. Collins, T. Meyer, Evolutionary origins of STIM1 and STIM2 within ancient Ca2+ signaling systems. Trends Cell Biol. 21, 202–211 (2011).

29. D. E. Schäffer, L. M. Iyer, A. M. Burroughs, L. Aravind, Functional innovation in the evolution of the calcium-dependent system of the eukaryotic endoplasmic reticulum. Front. Genet. 11, 34 (2020).

30. S. M. Emrich, R. E. Yoast, M. Trebak, Physiological functions of CRAC channels. Annu. Rev. Physiol. 84, 355–379 (2022).

31. P. G. Hogan, R. S. Lewis, A. Rao, Molecular basis of calcium signaling in lymphocytes: STIM and ORAI. Annu. Rev. Immunol. 28, 491–533 (2010).

32. M. Prakriya, R. S. Lewis, Store-operated calcium channels. Physiol. Rev. 95, 1383–1436 (2015).

33. J. W. Putney Jr, A model for receptor-regulated calcium entry. Cell Calcium 7, 1–12 (1986).

34. S. Feske, et al., A mutation in Orai1 causes immune deficiency by abrogating CRAC channel function. Nature 441, 179–185 (2006).

35. J. Liou, et al., STIM is a Ca2+ sensor essential for Ca2+-store-depletion-triggered Ca2+ influx. Curr. Biol. 15, 1235–1241 (2005).

36. J. Roos, et al., STIM1, an essential and conserved component of store-operated Ca2+ channel function. J. Cell Biol. 169, 435–445 (2005).

37. M. Vig, et al., CRACM1 is a plasma membrane protein essential for store-operated Ca2+ entry. Science 312, 1220–1223 (2006).

38. S. L. Zhang, et al., Genome-wide RNAi screen of Ca(2+) influx identifies genes that regulate Ca(2+) release-activated Ca(2+) channel activity. Proc. Natl. Acad. Sci. U. S. A. 103, 9357–9362 (2006).

39. C. Buro, et al., Transcriptome analyses of inhibitor-treated schistosome females provide evidence for cooperating Src-kinase and TGFβ receptor pathways controlling mitosis and eggshell formation. PLoS Pathog. 9, e1003448 (2013).

40. T. Quack, V. Wippersteg, C. G. Grevelding, Cell cultures for schistosomes - Chances of success or wishful thinking? Int. J. Parasitol. 40, 991–1002 (2010).

41. G. N. Huang, et al., STIM1 carboxyl-terminus activates native SOC, I(crac) and TRPC1 channels. Nat. Cell Biol. 8, 1003–1010 (2006).

42. R. E. Yoast, et al., The native ORAI channel trio underlies the diversity of Ca2+ signaling events. Nat. Commun. 11, 2444 (2020).

43. M. Trebak, G. S. J. Bird, R. R. McKay, J. W. Putney Jr, Comparison of human TRPC3 channels in receptor-activated and store-operated modes. Differential sensitivity to channel blockers suggests fundamental differences in channel composition. J. Biol. Chem. 277, 21617–21623 (2002).

44. X. Zhang, et al., Distinct pharmacological profiles of ORAI1, ORAI2, and ORAI3 channels. Cell Calcium 91, 102281 (2020).

45. B. A. McNally, M. Yamashita, A. Engh, M. Prakriya, Structural determinants of ion permeation in CRAC channels. Proc. Natl. Acad. Sci. U. S. A. 106, 22516–22521 (2009).

46. X. Hou, L. Pedi, M. M. Diver, S. B. Long, Crystal structure of the calcium release-activated calcium channel Orai. Science 338, 1308–1313 (2012).

47. A. Gudlur, et al., STIM1 triggers a gating rearrangement at the extracellular mouth of the ORAI1 channel. Nat. Commun. 5, 5164 (2014).

48. M. Prakriya, R. S. Lewis, Separation and characterization of currents through store-operated CRAC channels and Mg2+-inhibited cation (MIC) channels. J. Gen. Physiol. 119, 487–507 (2002).

49. A. Zweifach, R. S. Lewis, Rapid inactivation of depletion-activated calcium current (ICRAC) due to local calcium feedback. J. Gen. Physiol. 105, 209–226 (1995).

50. A. V. Yeromin, J. Roos, K. A. Stauderman, M. D. Cahalan, A store-operated calcium channel in Drosophila S2 cells. J. Gen. Physiol. 123, 167–182 (2004).

51. P. S.-W. Yeung, et al., Mapping the functional anatomy of Orai1 transmembrane domains for CRAC channel gating. Proc. Natl. Acad. Sci. U. S. A. 115, E5193–E5202 (2018).

52. I. Frischauf, et al., Transmembrane helix connectivity in Orai1 controls two gates for calcium-dependent transcription. Sci. Signal. 10 (2017).

53. R. Palty, C. Stanley, E. Y. Isacoff, Critical role for Orai1 C-terminal domain and TM4 in CRAC channel gating. Cell Res. 25, 963–980 (2015).

54. M. Prakriya, et al., Orai1 is an essential pore subunit of the CRAC channel. Nature 443, 230–233 (2006).

55. I. Frischauf, et al., Molecular determinants of the coupling between STIM1 and Orai channels: differential activation of Orai1-3 channels by a STIM1 coiled-coil mutant. J. Biol. Chem. 284, 21696–21706 (2009).

56. B. A. McNally, A. Somasundaram, M. Yamashita, M. Prakriya, Gated regulation of CRAC channel ion selectivity by STIM1. Nature 482, 241–245 (2012).

57. C. Y. Park, et al., STIM1 clusters and activates CRAC channels via direct binding of a cytosolic domain to Orai1. Cell 136, 876–890 (2009).

58. Y. Zhou, et al., Initial activation of STIM1, the regulator of store-operated calcium entry. Nat. Struct. Mol. Biol. 20, 973–981 (2013).

59. A. Gudlur, et al., Calcium sensing by the STIM1 ER-luminal domain. Nat. Commun. 9, 4536 (2018).

60. Y. Zhou, et al., The STIM1-binding site nexus remotely controls Orai1 channel gating. Nat. Commun. 7, 13725 (2016).

61. M. Fukushima, T. Tomita, A. Janoshazi, J. W. Putney, Alternative translation initiation gives rise to two isoforms of Orai1 with distinct plasma membrane mobilities. J. Cell Sci. 125, 4354–4361 (2012).

62. P. N. Desai, et al., Multiple types of calcium channels arising from alternative translation initiation of the Orai1 message. Sci. Signal. 8, ra74 (2015).

63. Y. Zhou, et al., STIM1 gates the store-operated calcium channel ORAI1 in vitro. Nat. Struct. Mol. Biol. 17, 112–116 (2010).

64. A. Lis, S. Zierler, C. Peinelt, A. Fleig, R. Penner, A single lysine in the N-terminal region of store-operated channels is critical for STIM1-mediated gating. J. Gen. Physiol. 136, 673–686 (2010).

65. B. A. McNally, A. Somasundaram, A. Jairaman, M. Yamashita, M. Prakriya, The C- and N-terminal STIM1 binding sites on Orai1 are required for both trapping and gating CRAC channels. J. Physiol. 591, 2833–2850 (2013).

66. H. Zheng, et al., Differential roles of the C and N termini of Orai1 protein in interacting with stromal interaction molecule 1 (STIM1) for Ca2+ release-activated Ca2+ (CRAC) channel activation. J. Biol. Chem. 288, 11263–11272 (2013).

67. I. Derler, et al., The extended transmembrane Orai1 N-terminal (ETON) region combines binding interface and gate for Orai1 activation by STIM1. J. Biol. Chem. 288, 29025–29034 (2013).

68. R. Palty, E. Y. Isacoff, Cooperative binding of stromal interaction molecule 1 (STIM1) to the N and C termini of calcium release-activated calcium modulator 1 (Orai1). J. Biol. Chem. 291, 334–341 (2016).

69. L. Niu, et al., STIM1 interacts with termini of Orai channels in a sequential manner. J. Cell Sci. 133, jcs239491 (2020).

70. K. A. Stauderman, CRAC channels as targets for drug discovery and development. Cell Calcium 74, 147–159 (2018).

71. J. Ishikawa, et al., A pyrazole derivative, YM-58483, potently inhibits store-operated sustained Ca2+ influx and IL-2 production in T lymphocytes. J. Immunol. 170, 4441–4449 (2003).

72. C. Zitt, et al., Potent inhibition of Ca2+ release-activated Ca2+ channels and T-lymphocyte activation by the pyrazole derivative BTP2. J. Biol. Chem. 279, 12427–12437 (2004).

73. S. Gao, et al., Mechanism of different spatial distributions of Caenorhabditis elegans and human STIM1 at resting state. Cell Calcium 45, 77–88 (2009).

74. K. M. Kim, et al., Distinct gating mechanism of SOC channel involving STIM-Orai coupling and an intramolecular interaction of Orai in Caenorhabditis elegans. Proc. Natl. Acad. Sci. U. S. A. 115, E4623–E4632 (2018).

75. C. Lorin-Nebel, J. Xing, X. Yan, K. Strange, CRAC channel activity in C. elegans is mediated by Orai1 and STIM1 homologues and is essential for ovulation and fertility. J. Physiol. 580, 67–85 (2007).

76. R. Qiu, R. S. Lewis, Structural features of STIM and Orai underlying store-operated calcium entry. Curr. Opin. Cell Biol. 57, 90–98 (2019).

77. P. S.-W. Yeung, M. Yamashita, M. Prakriya, Molecular basis of allosteric Orai1 channel activation by STIM1. J. Physiol. 598, 1707–1723 (2020).

78. J. H. Baraniak Jr, Y. Zhou, R. M. Nwokonko, D. L. Gill, The intricate coupling between STIM proteins and Orai channels. Curr. Opin. Physiol. 17, 106–114 (2020).

79. C. Humer, C. Romanin, C. Höglinger, Highlighting the multifaceted role of Orai1 N-terminal-and loop regions for proper CRAC channel functions. Cells 11, 371 (2022).

80. X. Hou, I. R. Outhwaite, L. Pedi, S. B. Long, Cryo-EM structure of the calcium release-activated calcium channel Orai in an open conformation. Elife 9 (2020).

81. N. Hirve, V. Rajanikanth, P. G. Hogan, A. Gudlur, Coiled-coil formation conveys a STIM1 signal from ER lumen to cytoplasm. Cell Rep. 22, 72–83 (2018).

